# Genomic instability caused by Arp2/3 complex inactivation results in micronucleus biogenesis and cellular senescence

**DOI:** 10.1101/2022.01.24.477450

**Authors:** Elena L. Haarer, Shirley Guo, Kenneth G. Campellone

## Abstract

The Arp2/3 complex is a ubiquitous actin nucleator with well-characterized activities in cell organization and movement, but its roles in chromatin-associated and cell cycle-related processes are relatively understudied. We investigated how the Arp2/3 complex affects genomic integrity, mitosis, and cell proliferation using mouse fibroblasts containing an inducible knockout (iKO) of the ArpC2 subunit. We show that permanent Arp2/3 ablation results in DNA damage, the formation of cytosolic micronuclei, and cellular senescence. Upon Arp2/3 depletion, cells undergo an abrupt proliferation arrest that is accompanied by activation of the tumor suppressor p53, upregulation of its downstream cell cycle inhibitor *Cdkn1a*/p21, and recognition of micronuclei by the cytosolic DNA sensor cGAS. Micronuclei arise in ArpC2 iKO cells due to chromosome segregation defects during mitosis and premature mitotic exits. Such phenotypes are explained by the presence of damaged chromatin fragments that fail to attach to the mitotic spindle, abnormalities in actin assembly during metaphase, and asymmetric microtubule architecture during anaphase. These studies establish functional requirements for the mammalian Arp2/3 complex in genome stability and mitotic spindle organization. They further expand our understanding of the intracellular mechanisms that lead to senescence and suggest that cytoskeletal dysfunction is an underlying factor in biological aging.

**Author Summary:** The actin cytoskeleton consists of protein polymers that assemble and disassemble to control the organization, shape, and movement of cells. However, relatively little is understood about how the actin cytoskeleton affects genome maintenance, cell multiplication, and biological aging. In this study, we show that knocking out the Arp2/3 complex, a core component of the actin assembly machinery, causes DNA damage, genomic instability, defects in chromosome partitioning during mitosis, and a permanent cell proliferation arrest called senescence. Since senescent cells are major contributors to both age-associated diseases and tumor suppression, our findings open new avenues of investigation into how natural or experimental alterations of cytoskeletal proteins impact the process of aging and the regulation of cancer.

## Introduction

The actin cytoskeleton consists of dynamic protein polymers that have well-known functions in cell morphogenesis and motility. Globular (G-) actin monomers are present in the cytoplasm and nucleus, and their polymerization into filamentous (F-) actin is driven by proteins called nucleators [1]. These include actin monomer-oligomerizing proteins, Formin-family nucleation/elongation proteins, and the Arp2/3 complex – a heteroheptameric actin assembly factor that binds to the sides of existing filaments and nucleates new filaments to create branched networks [2]. The Arp2/3 complex is highly conserved across almost all eukaryotes [3, 4] and is required for viability in such organisms; inactivation of genes encoding its subunits prevents growth of *S.cerevisiae* [5, 6] and *D.discoideum* [7] and is embryonic lethal in animals including *D.melanogaster* [8, 9], *C.elegans* [10, 11], and *M.musculus* [12–14]. However, the cellular basis underlying the essential nature of the Arp2/3 complex is not well understood.

Many processes that involve plasma membrane dynamics, especially cell adhesion and motility, rely on actin networks assembled by the Arp2/3 complex [15]. In fact, conditional knockouts in mice indicate that the complex is crucial for maintaining normal tissue architecture, promoting changes in cell shape, and powering cell migration during development [14,16–20].

These *in vivo* results are consistent with molecular and cellular studies of Arp2/3-mediated actin assembly using *in vitro* systems [21], dominant negative regulatory proteins [22], transient RNAi-mediated knockdowns [23], and pharmacological inhibitors of the complex [24, 25].

In contrast to the well-characterized roles of the Arp2/3 complex in protrusion and motility, its functions in nuclear processes are only beginning to emerge. During interphase, all 3 classes of actin nucleators promote nuclear actin filament assembly in response to acute DNA damaging agents [26–28]. In the nuclei of *Drosophila* and mammalian cells, Arp2/3-mediated actin polymerization is crucial for repositioning damaged heterochromatin to the nuclear periphery, which enables subsequent DNA repair activities [27, 28]. As a result, depletion of the Arp2/3 complex using RNAi in *Drosophila* larvae leads to chromosomal abnormalities and genomic instability [27]. Depletion of the Arp2/3 complex in human cells additionally results in defects in DNA damage-induced apoptosis [29].

Apart from their functions in chromatin-associated processes during interphase, actin and its nucleators, especially Formins and the Arp2/3 complex, are increasingly being found to support proper chromosome movements during meiosis and mitosis. In starfish oocytes, following nuclear envelope breakdown, several types of F-actin structures promote chromosome transport and coordinate capture by microtubules [30, 31]. Studies in mouse oocytes further indicate that actin filaments permeate the meiotic microtubule spindles and facilitate proper chromosome congression [32, 33]. Chemical inhibition of actin dynamics or genetic inactivation of Formin-2 prevents proper formation of kinetochore microtubules and leads to chromosome alignment and segregation errors [33, 34]. Similarly, during mitosis, several actin structures have been shown to interact with and possibly guide microtubule spindle components. Actin filaments that run between the microtubule spindle poles and F-actin fingers that project from the cell cortex into the spindle have been identified in *Xenopus* epithelial cells [35].

Centrosomes, which serve as major microtubule nucleation and organizing centers, are also sites of actin assembly [36]. The Arp2/3 complex localizes to centrosomes in multiple mammalian cell types, and pharmacological inhibition of Arp2/3 results in decreased centrosomal actin levels and impaired mitotic spindle formation [37–39]. Thus, disruption of either actin or Arp2/3 function during meiosis or mitosis can lead to defects in chromosome dynamics, highlighting the actin cytoskeleton as a key player in maintaining genomic integrity during nuclear division.

Although the effects of transient Arp2/3 depletion or inhibition on chromatin repair are now evident, and several aberrations in chromosome movement have been characterized, the impact of total and permanent Arp2/3 ablation on these processes has not been established. The development of several cellular systems for studying long-term Arp2/3 depletion or deletion has allowed more clear-cut assessments of the requirements for the Arp2/3 complex in a given cellular process [14,40–43]. For example, these models have already provided fundamental insights into the function of the Arp2/3 complex in cell migration. Studies using mouse embryonic fibroblasts (MEFs) expressing shRNAs targeting the ArpC2 and Arp2 subunits [40], embryonic stem cell-derived mouse fibroblasts lacking the ArpC3 subunit [14], and mouse fibroblasts harboring a tamoxifen-inducible knockout of the Arp3 subunit [43], indicate that the Arp2/3 complex is crucial for lamellipodia formation and directional migration. Additionally, mouse fibroblasts containing a tamoxifen-inducible knockout of the ArpC2 subunit exhibit impaired lamellipodia formation and reduced motility speeds [42].

To determine the outcome of Arp2/3 complex ablation on chromatin-associated processes related to cell viability and multiplication, we employed the inducible ArpC2 knockout cell model [42]. Our findings establish a key connection between Arp2/3 complex functions in genomic integrity during interphase and mitosis in normal cells to the biogenesis of micronuclei and induction of a cellular senescence pathway upon Arp2/3 inactivation.

## Results

### ArpC2 iKO cells undergo an abrupt proliferation arrest and morphological enlargement

Given that the Arp2/3 complex is required for viability in many eukaryotic organisms, we sought to better understand its essential nature – apart from its well-recognized roles in adhesion and motility – in mammalian cells. The Arp2/3 complex is composed of seven subunits: two <u>A</u>ctin-related <u>p</u>roteins (Arp2 and Arp3) and five <u>C</u>omplex subunits (ArpC1-C5), with ArpC2 and ArpC4 forming a structural core and multiple isoforms of Arp3, ArpC1, and ArpC5 providing peripheral diversity [4, 44]. Previous work indicates that MEFs subjected to RNAi-mediated depletion of the ArpC2 and Arp2 subunits are viable and remain culturable when generated in a genetic background lacking the *p16Ink4a/Arf* tumor suppressors [40]. More recently, to circumvent problems with knockdown instability, a conditional knockout model was created using *p16Ink4a/Arf* ^-/-^ mouse tail fibroblasts (MTFs) harboring a floxed *ArpC2* allele and engineered to express the CreER recombinase upon treatment with 4-hydroxytamoxifen (4-OHT) [42]. Since the latter system is inducible, causes a permanent loss of the critical ArpC2 subunit, and leads to degradation of other members of the complex, we adopted this Arp2/3 complex null cellular model for our studies. In all of our experiments, parental MTFs carrying the conditional *ArpC2* allele were exposed to DMSO to maintain a control (Flox) cell population or to 4-OHT to generate ArpC2 induced knockout (iKO) cells.

For initially assessing the kinetics of Arp2/3 complex depletion, DMSO- and 4-OHT-treated cells were collected at various timepoints, lysed, and immunoblotted with antibodies to the ArpC2 and Arp3 subunits plus antibodies to GAPDH and tubulin as loading controls (Fig 1A). After 1 day in 4-OHT, ArpC2 protein levels were reduced by nearly half compared to DMSO-treated Flox cells (Fig 1A and 1B). By 2 days, ArpC2 expression was diminished by approximately 80%, and a reduction in Arp3 levels became noticeable (Fig 1A and 1B). The amounts of both subunits continued to steadily decline in the iKO cells over time until they were absent following 5 days in 4-OHT (Fig 1A and 1B). DMSO and 4-OHT were removed from culture media after 6 days, but ArpC2 and Arp3 remained undetectable in the iKO population out to 13 days (Fig 1A and 1B). To independently confirm the loss of the Arp2/3 complex by fluorescence microscopy, Flox and iKO cells were stained with an antibody to label Arp3 and with fluorescent phalloidin to visualize F-actin. As expected, Flox cells exhibited prominent Arp3 staining within F-actin-rich peripheral membrane ruffles, whereas Arp3 staining and ruffles were both missing from the iKO cells (Fig 1C). Collectively, these results demonstrate that in this cellular context, the Arp2/3 complex knockout is rapid, complete, and stable over time.

**Fig 1.**
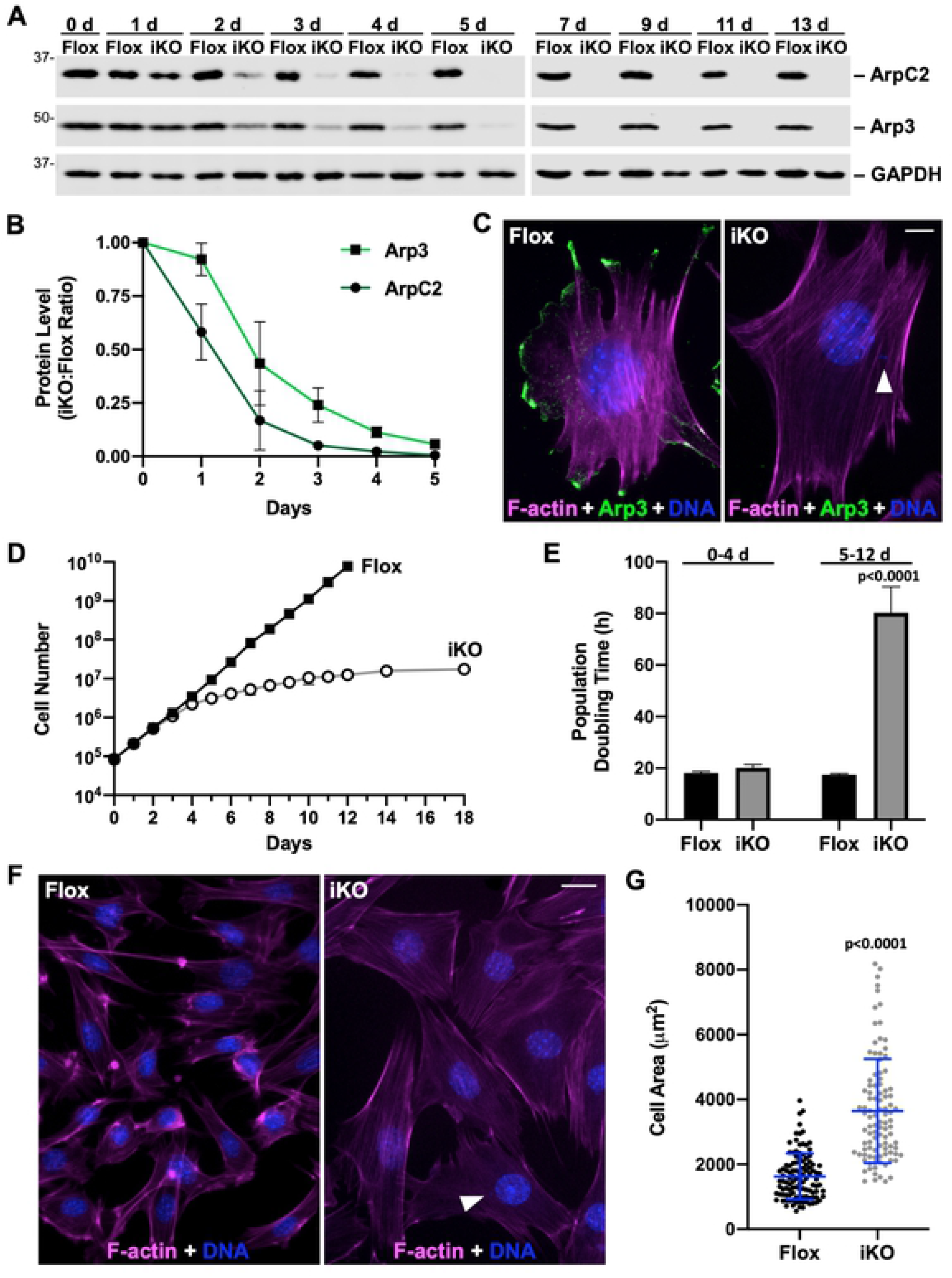
ArpC2 iKO cells undergo a proliferation arrest and enlargement. **(A)** Mouse fibroblasts were treated with DMSO (Flox) or 4-OHT (iKO) for 0-6d and collected at 0-13d. Samples were lysed, subjected to SDS-PAGE, and immunoblotted with antibodies to ArpC2, Arp3, and GAPDH. **(B)** ArpC2 and Arp3 band intensities were normalized to GAPDH or tubulin band intensities and plotted as the iKO:Flox ratio. Each dot represents the mean ratio ±SD from 2-3 experiments. **(C)** Flox and iKO cells were fixed at 7d and stained with phalloidin (F-actin; magenta), an Arp3 antibody (green), and DAPI (DNA; blue). The arrowhead highlights a micronucleus. Scale bar, 10µm. **(D)** Flox and iKO cell titers were quantified at 0-18d. Each point represents the mean ±SD from 2 Flox and 2-3 iKO experiments, except for the 14d and 18d timepoints, which were from a representative iKO sample that did not exhibit outgrowth of colonies expressing ArpC2. **(E)** Flox and iKO population doubling times were quantified daily from 0-4d and 5-12d. For each time range, the bar represents the mean doubling time ±SD from n=3 experiments. **(F)** Flox and iKO cells were fixed at 7d and stained with phalloidin and DAPI. The arrowhead highlights a micronucleus. Scale bar, 25µm. **(G)** Flox and iKO cells were outlined in ImageJ and their areas quantified. Each dot represents an individual cell and the lines represent the mean area ±SD from analyses of 100 cells.

To determine the impact of abolishing Arp2/3 complex expression on cell proliferation, we next quantified cell titers on a daily basis following the addition of DMSO or 4-OHT. For the first 3 days, Flox and iKO cells multiplied at identical rates, but by 4 days, the growth characteristics of Flox and iKO cultures began to diverge, and at 5 days, the iKO samples were proliferating at a clearly slower pace (Fig 1D). After approximately 10-12 days, virtually all iKO cells stopped dividing (Fig 1D). To quantify the differences in cell multiplication rates, we calculated the population doubling times in the 0-4 and 5-12 day time periods following DMSO or 4-OHT exposure. While the doubling times were similar for Flox and iKO populations (18h vs. 20h) during the first interval, the iKO doubling times quadrupled to >80h in the 5-12 day range (Fig 1E). Moreover, cell counts in iKO samples remained unchanged for an additional week (Fig 1D), except in instances where colonies of 4-OHT escapees or revertants re-expressing the Arp2/3 complex emerged (not depicted). Consistent with a requirement for the Arp2/3 complex in intrinsic apoptosis [29], apoptotic cell phenotypes were not observed in the iKO cell population. Thus, following loss of the Arp2/3 complex, MTF cells undergo an abrupt and stable proliferation arrest.

When examining the different growth characteristics of the Flox and iKO cultures, it also became apparent that the two cell types had distinct morphologies and sizes. Fluorescent phalloidin staining, in addition to revealing a lack of F-actin-rich ruffles (Fig 1C), demonstrated that iKO cells were flatter and larger than Flox cells (Fig 1F). Quantification of Flox and iKO cell areas showed that the iKO cells were, on average, about twice as large as Flox cells (Fig 1G). So in addition to losing their ability to multiply, Arp2/3-deficient cells display significant increases in their size.

### ArpC2 iKO cells exhibit the canonical nuclear and cytoplasmic features of senescence

A loss of proliferative capacity and an increase in adherent cell area are common characteristics of senescent cells. Cellular senescence refers to a permanent state of replicative arrest [45, 46], and is reflected in several additional physiological changes, including increased production of pro-inflammatory proteins, a response known as the senescence-associated secretory phenotype, or SASP [47, 48]. Notably, previous global gene expression profiling using the ArpC2/Arp2-depleted MEF model revealed that several genes common to the SASP were up- regulated [41], suggesting a link between the loss of Arp2/3 function and this aspect of cellular senescence. To investigate whether the ArpC2 iKO cells also displayed this senescence feature, we performed RT-PCR (Fig 2A) and RT-qPCR (Fig 2B) to compare transcript levels for Interleukin-6 (Il-6), a pro-inflammatory cytokine consistently present in the SASPs derived from senescent cells with diverse origins [49, 50]. In agreement with findings from the ArpC2/Arp2 RNAi MEF studies, *Il-6* expression was greater in iKO cells than in Flox cells at 3, 6, and 9 days after the onset of 4-OHT treatment (Fig 2A). RT-qPCR revealed that *Il-6* transcript levels were nearly 4-fold higher in the iKO compared to Flox cells at 9 days (Fig 2B), indicating that a permanent loss of the Arp2/3 complex induces the production of this key SASP component.

**Fig 2.**
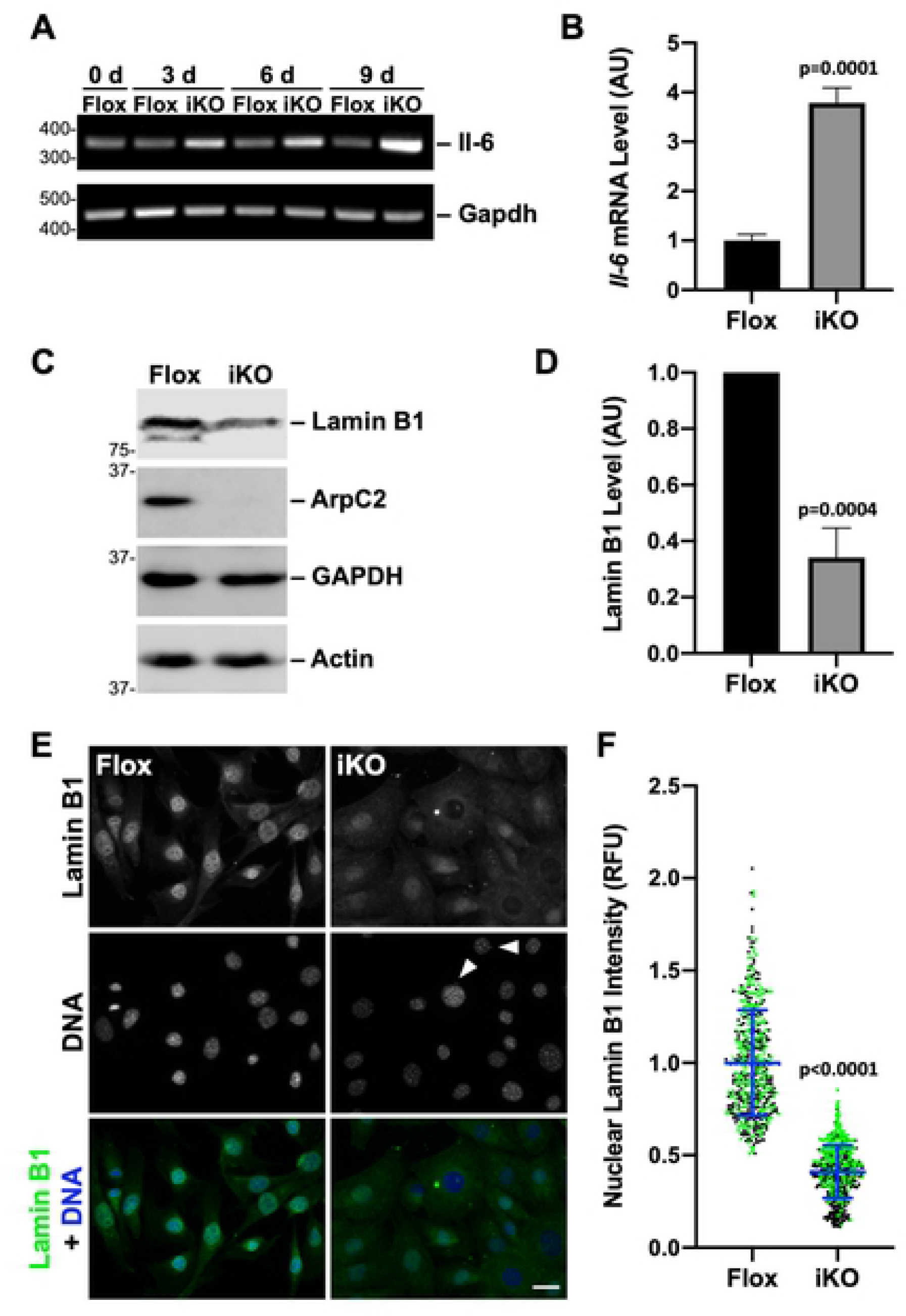
Loss of the Arp2/3 complex results in elevated *Il-6* transcript and decreased nuclear Lamin B1 levels. **(A)** Mouse fibroblasts were treated with DMSO (Flox) or 4-OHT (iKO) for 0-6d and collected at 0, 3, 6, and 9d. RNA was isolated and RT-PCR performed using primers for *Il-6* and *Gapdh*. PCR products were visualized on ethidium bromide stained agarose gels. **(B)** RT-qPCR was performed using primers for *Il-6* and *Gapdh* at 9d. *Il-6* product levels were normalized to *Gapdh*. Each bar represents the mean transcript abundance ±SD from n=3 experiments. AU = Arbitrary Units. **(C)** Flox and iKO cells were collected at 10d and immunoblotted with antibodies to Lamin B1, ArpC2, GAPDH, and actin. **(D)** Lamin B1 band intensity was normalized to GAPDH and actin band intensities. Each bar represents the mean intensity ±SD from n=3 experiments. **(E)** Flox and iKO cells were fixed at 9d and stained with a Lamin B1 antibody (green) and DAPI (DNA; blue). Arrowheads point to micronuclei. Scale bar, 25µm. **(F)** Nuclear Lamin B1 levels were quantified by outlining the DAPI-stained nucleus of each cell in ImageJ and measuring the mean Lamin B1 pixel intensity. Each dot represents an individual cell and the line depicts the average Lamin B1 pixel intensity from analyses of 492- 556 cells pooled from n=2 experiments (denoted in black or green). RFU = Relative Fluorescence Units.

Senescence is also frequently associated with changes in nuclear structure, including decreased levels of Lamin B1, a structural protein of the nuclear lamina [51]. To determine if Lamin B1 abundance was altered by the deletion of ArpC2, we immunoblotted Flox and iKO cells for Lamin B1 and found that Lamin B1 protein levels were indeed 3-fold lower in iKO cell populations (Fig 2C and 2D). To confirm the reduction in Lamin B1 expression, Flox and iKO cells were also subjected to immunofluorescence microscopy. In Flox cells, nuclear Lamin B1 staining was consistently bright (Fig 2E and 2F). Contrastingly, in iKO cells, nuclear Lamin B1 levels were visibly lower (Fig 2E and 2F). In agreement with the immunoblotting data, quantification of nuclear fluorescence intensities demonstrated that, on average, Lamin B1 was nearly 3-fold less abundant in iKO cells (Fig 2F).

In addition to the production of a transcriptional SASP response and a decrease in their nuclear Lamin B1 levels, senescent cells typically exhibit a cytoplasmic senescence-associated β-galactosidase (SA-βgal) activity at pH 6 [52]. This is the most widely accepted biomarker of senescence. SA-βgal staining in Flox and iKO cells from 0-22 days revealed that by 7 days, and at later timepoints, the number of SA-βgal-positive cells was significantly higher in iKO than Flox populations (Fig 3A and 3B). In many instances, SA-βgal activity is thought to reflect an increase in lysosomal content [53, 54]. So to assess lysosomal abundance, we treated Flox and iKO cells with LysoTracker, a fluorescent probe that labels acidic intracellular structures, and examined the cells microscopically. LysoTracker intensely stained discrete circular puncta resembling lysosomes in Flox cells, but broadly stained large portions of the cytoplasm in iKO cells (Fig 3C). Quantification of LysoTracker fluorescence at day 9 revealed that more than

**Fig 3.**
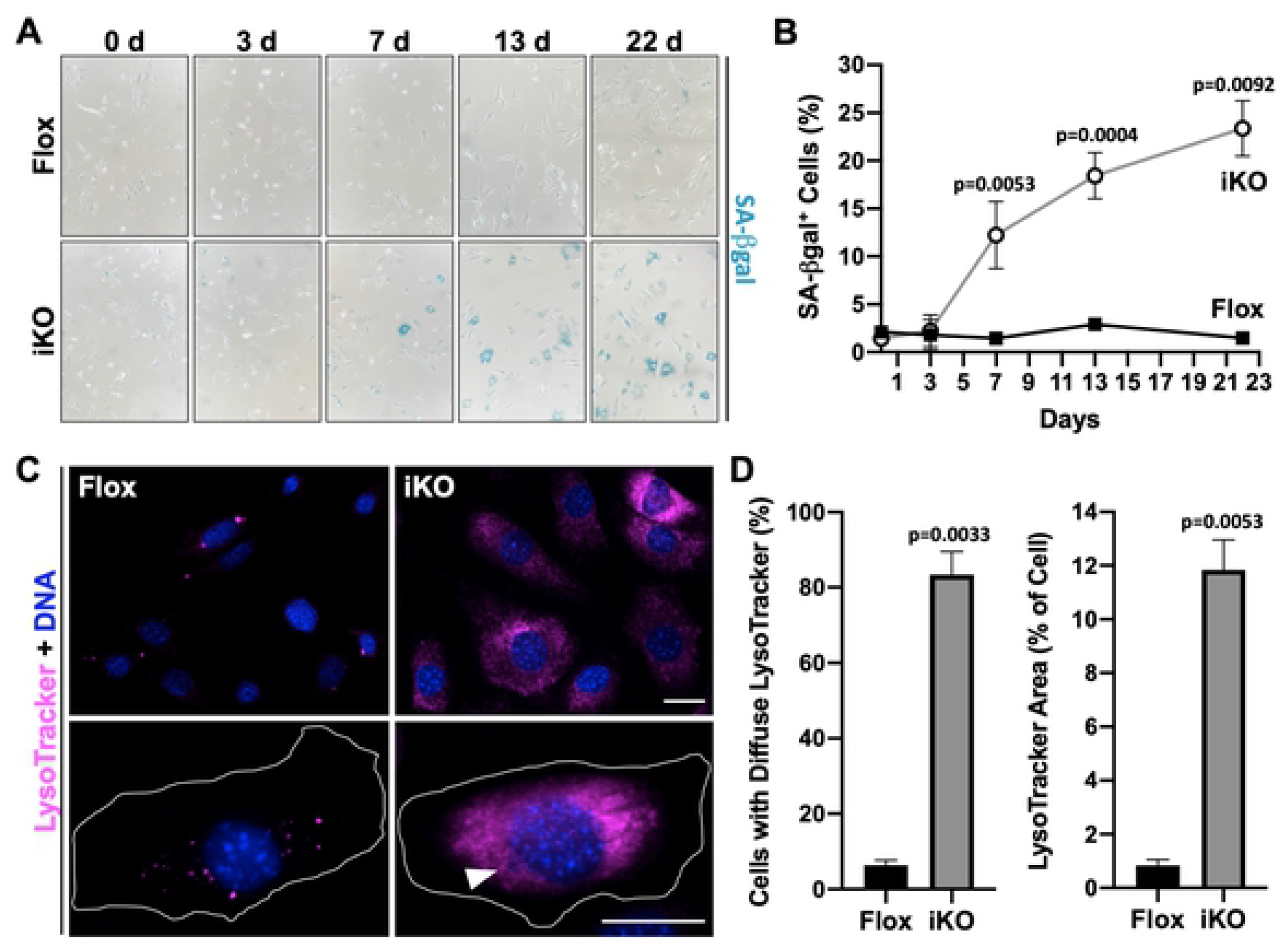
SA-βgal activity and lysosomal content are increased in ArpC2 iKO cells. **(A)** Mouse fibroblasts were treated with DMSO (Flox) or 4-OHT (iKO) for 0-6d and senescence-associated beta-galactosidase (SA-βgal) staining was performed over a 22d period. **(B)** The % of SA-βgal-positive cells was quantified by calculating the number of intensely blue-colored cells divided by the total number of cells. Each point represents the mean % ±SD from a representative 0d timepoint, n=2 experiments for the 3d and 22d timepoints, and n=3 experiments for the 7d and 13d timepoints (727-2197 cells per point). **(C)** Flox and iKO cells at 9d were treated with LysoTracker (magenta), fixed, and stained with DAPI (DNA; blue). The magnifications illustrate punctate versus diffuse LysoTracker staining, and the arrowhead indicates the position of a micronucleus. Scale bars, 25µm. **(D)** The % of cells exhibiting diffuse LysoTracker staining was quantified by scoring the number of cells with broad instead of punctate cytoplasmic fluorescence and dividing by the total number of cells. The relative area within each cell occupied by LysoTracker was quantified using the threshold function in ImageJ. Each bar represents the mean % ±SD from n=2 experiments (151-155 cells per bar).

80% of iKO cells versus 5% of Flox cells showed the more expansive staining pattern (Fig 3D). Furthermore, >10% of the areas within iKO cells stained positive for LysoTracker compared to <1% of the areas in Flox cells (Fig 3D). Thus, the arrest, morphological, and SASP observations, when taken together with these Lamin B1, SA-βgal, and LysoTracker staining results, establish that ArpC2 iKO cells undergo senescence.

### The formation of micronuclei and DNA damage clusters precedes iKO cell senescence

Cellular senescence can be induced by a variety of stimuli, including DNA damage, oncogene activation, telomere shortening, and mitochondrial dysfunction [55–57]. Micronuclei and cytosolic chromatin fragments are common senescence-associated phenotypic traits and influence the senescent state [58–60]. During the course of our characterization of the senescence parameters described above, we noticed that iKO cells frequently contained micronuclei whereas Flox cells did not (Fig 1C; Fig 2E; Fig 3C). Following specific examinations of samples containing DAPI-stained DNA, it became obvious that small cytosolic micronuclei were often present in iKO cells (Fig 4A). Of iKO cells with micronuclei, most had one micronucleus (Fig 4B; i, ii, iii), but some harbored 2 or 3 micronuclei (Fig 4B; iv, v, vi).

**Fig 4.**
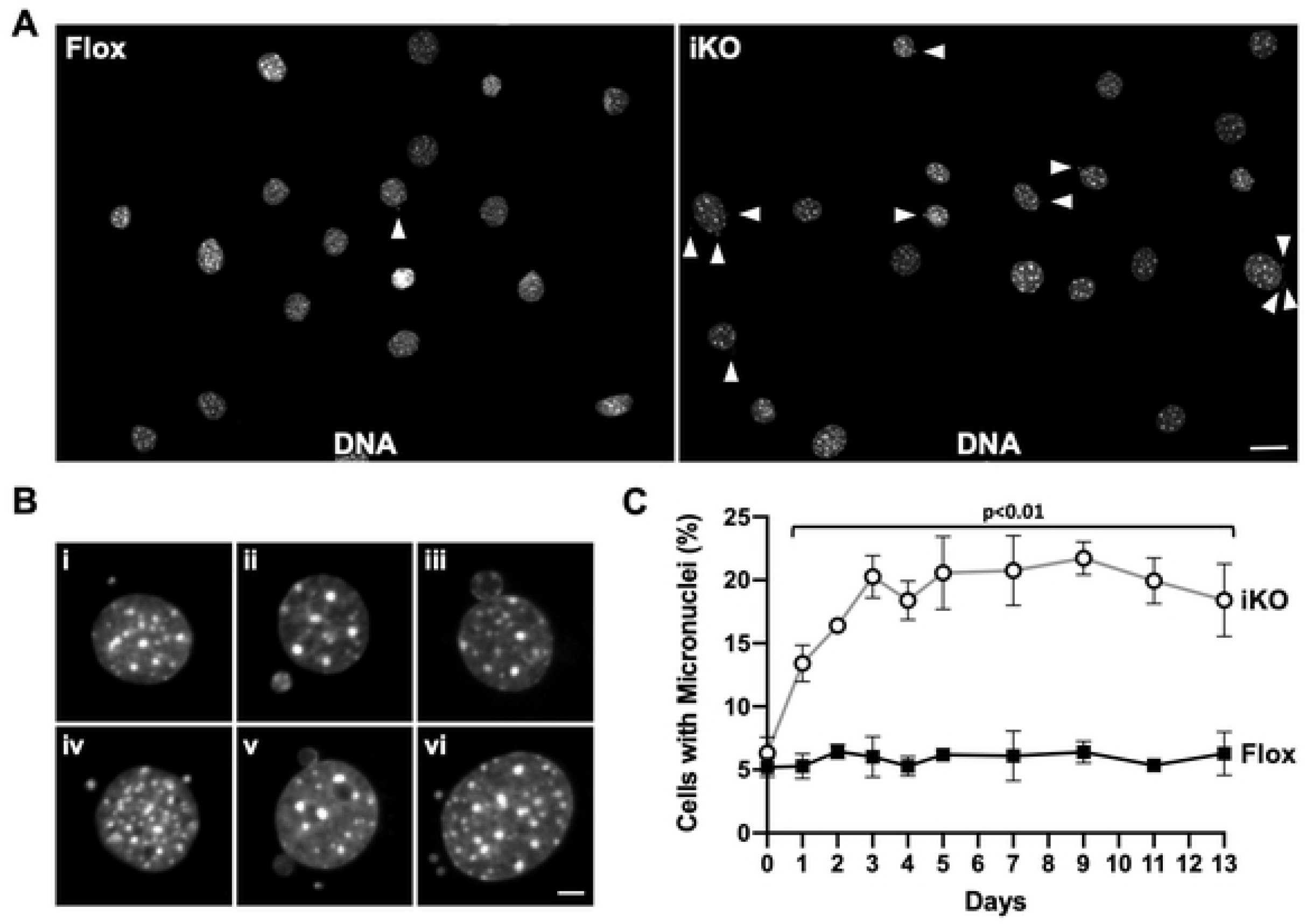
Arp2/3 complex depletion leads to the formation of cytoplasmic micronuclei. **(A)** Mouse fibroblasts were treated with DMSO (Flox) or 4-OHT (iKO) for 6d, fixed at 7d, and stained with DAPI. Arrowheads point to micronuclei. Scale bar, 25µm. **(B)** Magnifications show different micronucleus phenotypes in iKO cells. Scale bar, 5µm. **(C)** The % of cells with micronuclei was quantified over a 13d period following DMSO or 4-OHT exposure. Each point represents the mean % ±SD from n=3 experiments, except for the 2d and 4d timepoints, which are from n=2 experiments (432-631 cells per point in each experiment).

Micronuclei ranged in size and could be found completely detached (Fig 4B; i, ii, vi) or tethered to the periphery (Fig 4B; iii, iv, v) of the main nucleus. We next quantified the timing and frequency with which micronuclei formed in Flox and iKO cells. Surprisingly, after just one day in 4-OHT, about 13% of iKO cells had micronuclei (Fig 4C). The percentage of iKO cells with micronuclei plateaued at approximately 20% by 3 days and remained steady out to 13 days, whereas the proportion of Flox cells with micronuclei stayed at around 5% throughout the entire time course (Fig 4C). Therefore, a notably rapid formation of micronuclei precedes the proliferation arrest and SA-βgal positivity that results from *ArpC2* inactivation.

Since micronuclei are often indicative of genomic instability, we next assessed the extent of DNA damage in Flox and iKO cells. Staining with an antibody to the modified histone protein H2AX (γH2AX), which is phosphorylated in response to double-stranded DNA breaks [61], demonstrated that iKO cells contained prominent DNA damage clusters in their nuclei and micronuclei (Fig 5A). Upon closer inspection, Flox cells typically exhibited no or diffuse γH2AX staining in their nuclei (Fig 5B), whereas iKO cells frequently had 1-3 isolated γH2AX clusters that localized to the nuclear periphery and/or within micronuclei (Fig 5B). Quantification of the percentage of cells with γH2AX clusters following the addition of DMSO or 4-OHT revealed that clusters began to increase in iKO cells by 2 days, and that from 3 days onward, clusters were significantly more common in iKO cells than in Flox cells (Fig 5C). The fraction of iKO cells with γH2AX clusters leveled out at approximately 25%, while the proportion of Flox cells containing clusters was always 3-6% (Fig 5C). These results are consistent with previous observations in which *Drosophila* and mouse cells exposed to ionizing radiation and subjected to RNAi-mediated Arp2/3 depletion were found to contain DNA damage and micronuclei [27]. However, our results indicate that even without exposure to acute genotoxic agents, losing the Arp2/3 complex can cause an accumulation of damaged DNA elements that incorporate into micronuclei, thereby illustrating that the Arp2/3 complex is a crucial player in maintaining genomic integrity under relatively normal cell culture conditions.

**Fig 5.**
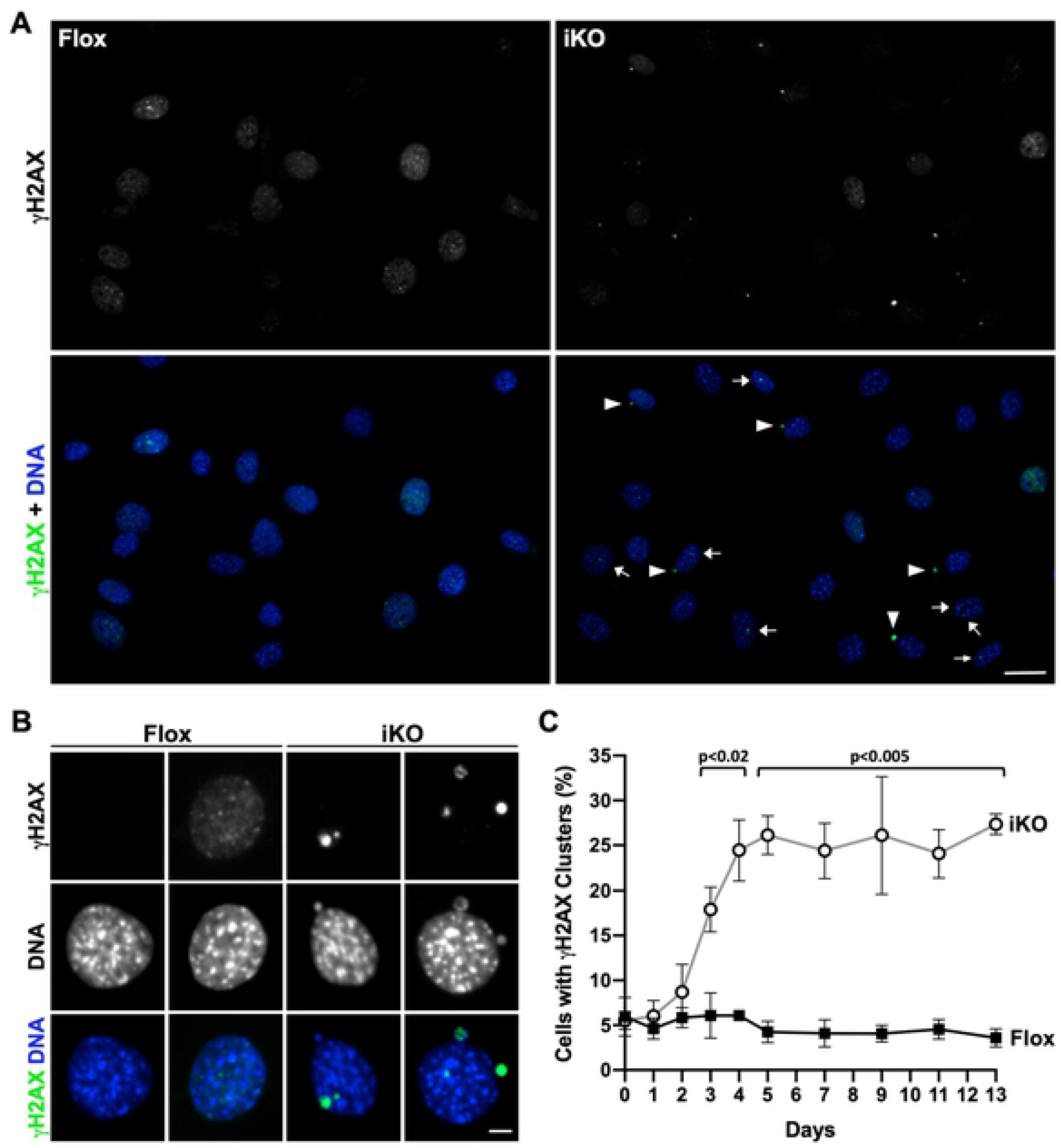
Prominent DNA damage clusters are found in the nuclei and micronuclei of ArpC2 iKO cells. **(A)** Mouse fibroblasts were treated with DMSO (Flox) or 4-OHT (iKO) for 6d, fixed at 7d, and stained with a γH2AX antibody (green) and DAPI (DNA; blue). Arrowheads point to γH2AX clusters in micronuclei and arrows indicate γH2AX clusters in nuclei. Scale bar, 25µm. **(B)** Magnifications show no or diffuse γH2AX staining in Flox nuclei and γH2AX clusters in iKO cell nuclei and micronuclei. Scale bar, 5µm. **(C)** The % of cells with γH2AX clusters was quantified over a 13d period following DMSO or 4-OHT exposure. Each point represents the mean % ±SD from n=3 experiments, except for 2d and 4d timepoints, which are from n=2 experiments (432-631 cells per point in each experiment).

### Activation of p53 and upregulation of *Cdkn1a*/p21 accompany the ArpC2 iKO cell arrest

In light of the biogenesis of γH2AX clusters and micronuclei, we hypothesized that a DNA damage response might be activated in the ArpC2 iKO cells. To explore this possibility, we immunoblotted Flox and iKO cell extracts with antibodies to the crucial tumor suppressor protein and transcription factor p53, which typically becomes stabilized, phosphorylated, and enriched in the nucleus following DNA damage [62, 63]. Consistent with this expectation, by 3 days after 4-OHT vs DMSO treatment, p53 levels were higher in iKO than in Flox cells, and p53 was phosphorylated on Serine-15 (Fig 6A). Furthermore, immunofluorescence microscopy revealed that nuclear p53 fluorescence was more intense in iKO cells than in Flox cells (Fig 6B). These results imply that a p53-mediated DNA damage response is induced in Arp2/3-depleted cells.

**Fig 6.**
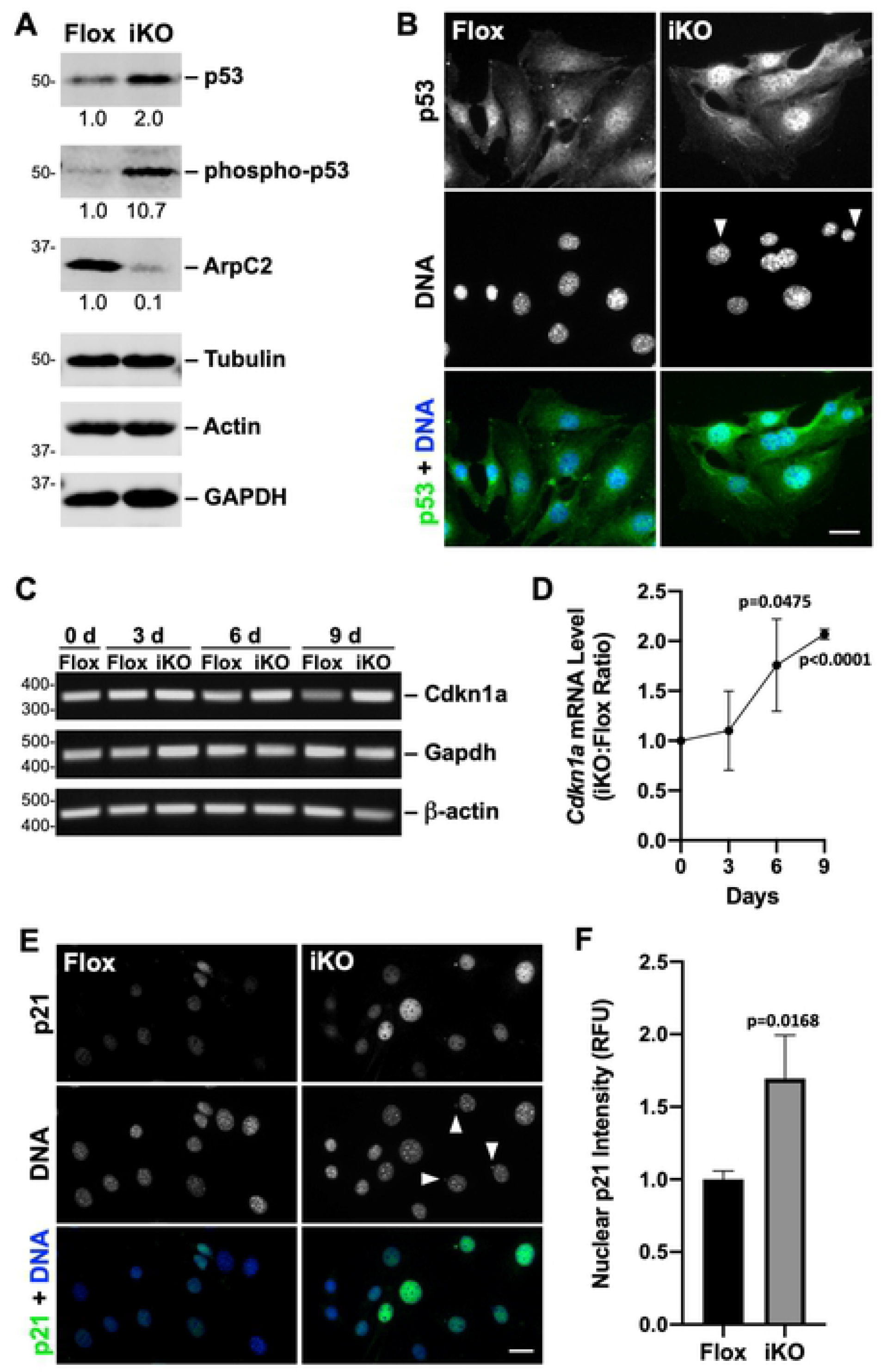
Activation of p53 and upregulation of *Cdkn1a*/p21 occur in ArpC2 iKO cells. **(A)** Mouse fibroblasts were treated with DMSO (Flox) or 4-OHT (iKO) for 3d, collected, and immunoblotted with antibodies to p53, phospho-p53(Ser15), ArpC2, tubulin, actin, and GAPDH. Total and phosphorylated p53 intensities relative to loading controls are indicated beneath the respective bands. **(B)** Flox and iKO cells were fixed at 3d and stained with a p53 antibody (green) and DAPI (DNA; blue). Arrowheads point to micronuclei. Scale bars, 25µm. **(C)** Flox and iKO cells were collected at 0, 3, 6, and 9d. RNA was isolated and RT-PCR performed using primers for *Cdkn1a*, *Gapdh* and *β-actin*. PCR products were visualized on ethidium bromide stained agarose gels. **(D)** RT-qPCR was performed using primers for *Cdkn1a* and *Gapdh* at 0, 3, 6, and 9d. *Cdkn1a* product levels were normalized to *Gapdh* and plotted as the iKO:Flox ratio. Each point represents the mean ratio ±SD from n=3 experiments. **(E)** Flox and iKO cells were fixed at 6d and stained with a p21 antibody (green) and DAPI. **(F)** Nuclear p21 fluorescence was quantified by outlining the DAPI-stained nucleus of each cell in ImageJ and measuring the mean p21 pixel intensity. Each bar represents the mean intensity ±SD from n=3 experiments (653-658 cells per bar). RFU = Relative Fluorescence Units.

Two of the major factors involved in the cell cycle arrest that leads to senescence are the cyclin-dependent kinase inhibitors *Cdkn2a*/p16INK4A (p16) and *Cdkn1a*/p21CIP/WAF (p21). Elevated levels of both of these anti-proliferative transcripts/proteins are frequently used as indicators of the senescent state [55, 64], although populations of cells expressing high levels of p16 appear to be distinct from those expressing high levels of p21, at least in tissues from aged mice [65]. Since p21 is a known transcriptional target of p53 [66, 67], and the MTFs used in our studies lack p16, we postulated that *Cdkn1a*/p21 was associated with the onset of senescence in the iKO cells. To test this, we performed RT-PCR (Fig 6C) and RT-qPCR (Fig 6D) to compare *Cdkn1a* transcript levels in Flox and iKO cells over a 9-day time period. Indeed, *Cdkn1a* expression appeared greater in the iKO cells than in the Flox cells at 6 and 9 days after the initiation of 4-OHT treatment (Fig 6C). RT-qPCR revealed that *Cdkn1a* transcript levels were doubled in the iKO cells relative to the Flox cells at 9 days (Fig 6D), showing that deletion of the Arp2/3 complex leads to an upregulation of this key cell cycle regulator. To evaluate whether the increase in *Cdkn1a* transcript corresponded to greater p21 protein levels, Flox and iKO cells were treated with antibodies to p21 and subjected to immunofluorescence microscopy (Fig 6E). Quantification of p21 nuclear intensity at the 6 day timepoint verified that p21 levels were significantly higher in iKO than in Flox cells (Fig 6E and 6F). Together, the p53 enrichment and p21 upregulation in iKO cells support the idea that a DNA damage response signaling pathway accompanies cell cycle arrest after the Arp2/3 complex is removed.

### Cytoplasmic cGAS and STING are recruited to micronuclei

In addition to the above nuclear changes that took place upon Arp2/3 ablation, it seemed likely that cytoplasmic changes arising from the presence of micronuclei in ArpC2 iKO cells would also be linked to the onset of senescence. We hypothesized that a cytosolic DNA detection and signaling pathway involving the cyclic GMP-AMP Synthase (cGAS) enzyme, which recognizes extra-nuclear chromatin and relays a signal to the downstream effector molecule STING [60, 68,69], might also be activated in iKO cells. Tagged cGAS can be recruited to micronuclei, and through its detection of cytosolic DNA and activation of STING, promotes pro-senescence and pro-inflammatory gene expression [70–72]. To determine if tagged cGAS localizes to the micronuclei in ArpC2 iKO cells, we transiently transfected MTFs with plasmids encoding mCherry-cGAS or mCherry as a control (Fig 7A), and treated them with 4-OHT to induce *ArpC2* deletion. mCherry was highly expressed in the iKO cells but was not recruited to micronuclei (Fig 7B). In contrast, mCherry-cGAS showed intense localization to micronuclei (Fig 7B), indicating that it can detect the cytosolic DNA in iKO cells.

**Fig 7.**
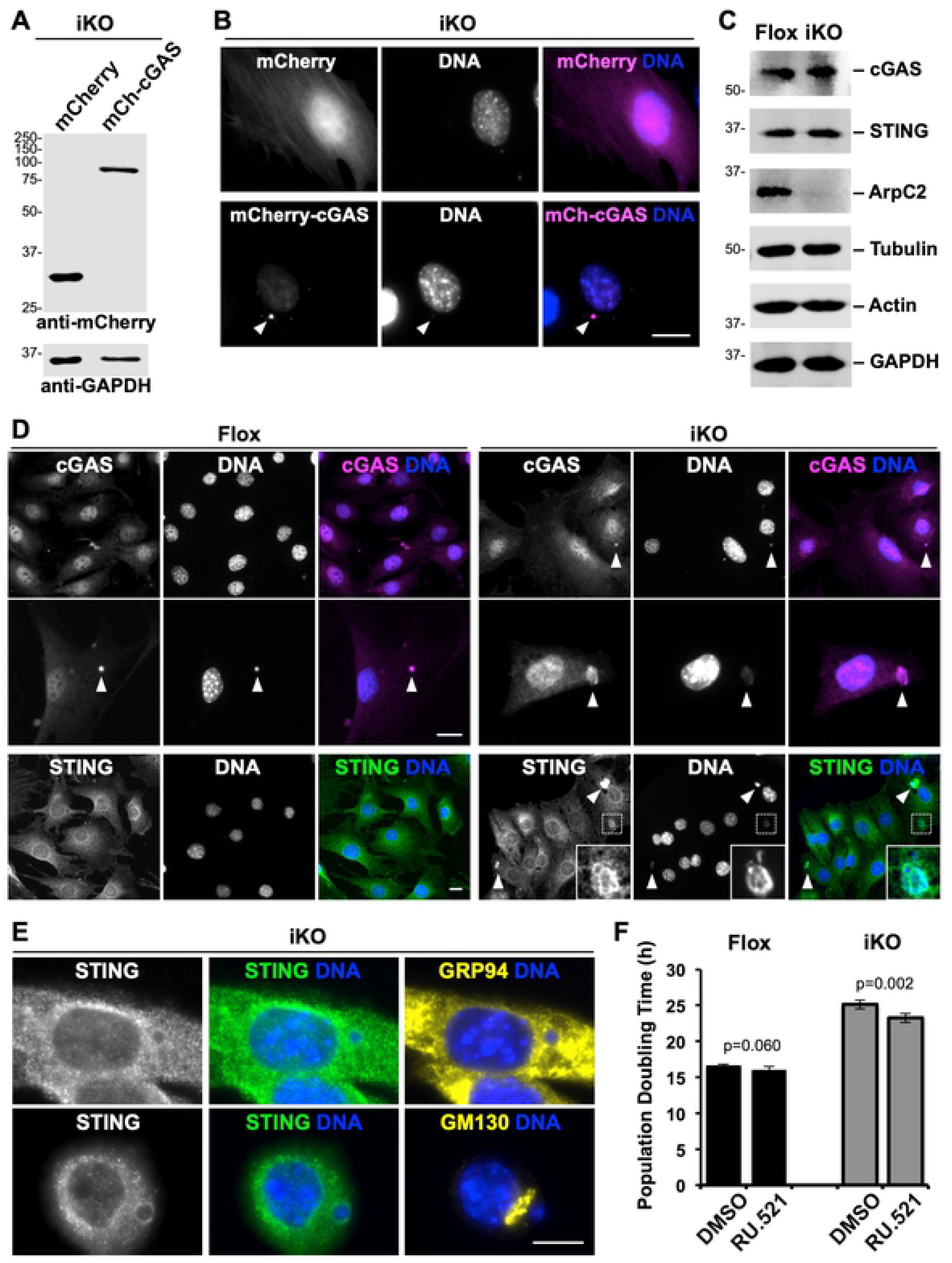
cGAS and STING are recruited to micronuclei. **(A)** Mouse fibroblasts were treated with 4-OHT (iKO), transfected with plasmids encoding mCherry or mCherry-cGAS, collected, and immunoblotted with antibodies to mCherry and GAPDH. **(B)** iKO cells expressing mCherry or mCherry-cGAS (magenta) were fixed at 1-2d and stained with DAPI (DNA; blue). Arrowheads point to a cGAS-positive micronucleus. Scale bars, 20µm. **(C)** Cells were treated with DMSO (Flox) or 4-OHT (iKO) for 3-4d and immunoblotted with antibodies to cGAS, STING, ArpC2, tubulin, actin, and GAPDH. **(D)** Flox and iKO cells were fixed at 3d and stained with cGAS (magenta) or STING (green) antibodies and DAPI. Arrowheads point to cGAS- or STING- positive micronuclei. 38.7% of micronuclei-containing iKO cells were cGAS-positive (75 cells from n=2 experiments). Insets show STING localization around an iKO micronucleus. **(E)** iKO cells were fixed at 3d and stained with STING antibodies (green) and either a GRP94 or GM130 antibody (yellow). **(F)** Cells were treated with DMSO or 4-OHT for 3d, switched to media containing DMSO or RU.521, and population doubling times were quantified daily from 4-6d. Each bar represents the mean doubling time ±SD from n=5 experiments.

We next wanted to determine the localization of endogenous cGAS and STING in the MTFs. Immunoblotting indicated that cGAS and STING were expressed at similar levels in Flox and iKO cells (Fig 7C), so to test whether cGAS and/or STING were recruited to the micronuclei, we visualized these proteins via immunofluorescence microscopy after 3-4 days, when micronuclei become more abundant in iKO cells and just before Flox and iKO multiplication rates begin to diverge. Consistent with previous studies [73, 74], endogenous cGAS was present in the nuclei of both Flox and iKO cells (Fig 7D). It was also readily apparent in nearly 40% of iKO cell micronuclei (Fig 7D). The rare micronuclei that formed in Flox cells also recruited cGAS (Fig 7D magnification). For STING, an integral membrane protein that localizes to organelles of the conventional secretory pathway, including the endoplasmic reticulum (ER) and Golgi [75], immunofluorescence revealed a speckled ER-like localization in both Flox and iKO cells, and enrichment around micronuclei (Fig 7D inset). Immunolabeling for STING, the ER chaperone GRP94, and the *cis*-Golgi protein GM130 implied that ER and not Golgi membranes are more likely to surround subsets of micronuclei in iKO cells (Fig 7E). These findings are consistent with a function for cGAS in recognizing damaged DNA in the cytosol and initiating a signaling pathway that locally activates STING near micronuclei in Arp2/3-deficient cells.

To explore whether cGAS affects the cell proliferation arrest that takes place when the Arp2/3 complex is lost, MTFs were exposed to 4-OHT for 3 days to induce the deletion of *ArpC2* and then transferred into media containing the cGAS inhibitor RU.521 [76] or DMSO as a control. Quantification of cell replication rates during days 4-6 following the onset of 4-OHT treatment revealed that RU.521 caused a modest but statistically faster population doubling time in the iKO cells (Fig 7F), implying that cGAS inhibition has the potential to oppose the initiation of senescence in some of the cells in this experimental system. Collectively, the above localization and inhibitor results support the idea that cGAS/STING signaling contributes to the establishment of senescence in Arp2/3-deficient cells.

### Micronuclei form as a result of mitotic defects in ArpC2 iKO cells

Because ArpC2 iKO cells accumulate DNA damage, form micronuclei with high frequency, and undergo senescence in a p53/p21- and cGAS/STING-associated manner, we next wanted to determine how this critical process of micronucleus biogenesis takes place. Micronuclei can form as a result of chromosome missegregation during mitosis and from expulsion of damaged DNA from the nucleus during interphase [58,77–80]. To differentiate between the possibilities that micronuclei are a product of defects in mitosis versus aberrant nuclear remodeling in interphase, we expressed the GFP-tagged histone H2B in MTFs and visualized chromatin dynamics in live cells. These experiments were performed within the first 2 days of DMSO or 4-OHT treatment, when Arp2/3 complex levels were declining the fastest (Fig 1B), the cells were still dividing rapidly (Fig 1E), and the incidence of micronucleus formation was highest (Fig 4C).

Timelapse imaging of H2B-GFP-expressing Flox cells revealed that the majority of mitoses resulted in an equal partitioning of nuclear chromatin into two daughter cells (Fig 8A).

**Fig 8.**
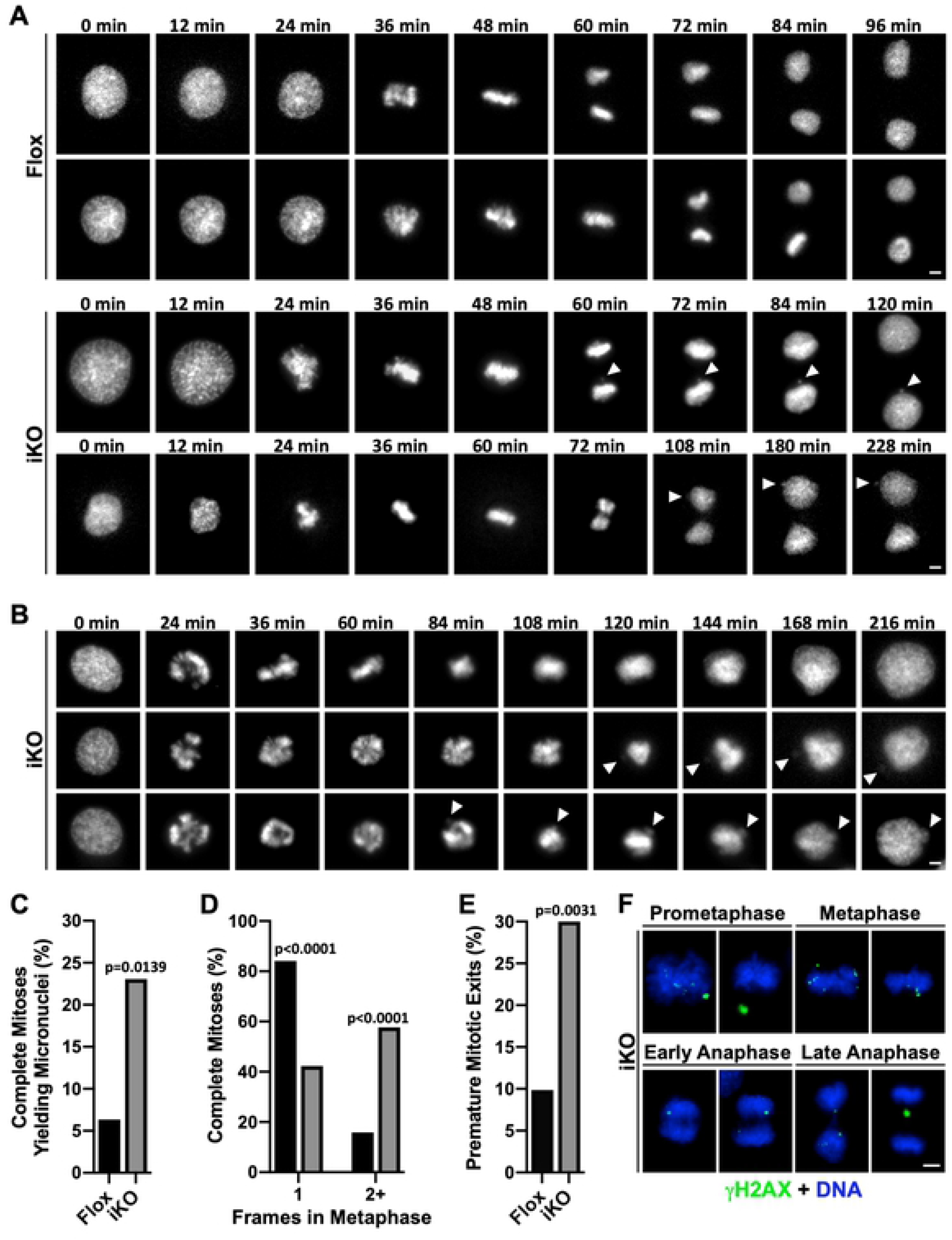
Micronuclei form due to chromatin segregation errors and premature mitotic exits in ArpC2 iKO cells. (A-B) Live H2B-GFP-expressing mouse fibroblasts were imaged every 12min from 1-2d following DMSO (Flox) or 4-OHT (iKO) exposure. Panel A shows completed mitoses, while panel B shows iKO cells exiting mitosis prematurely. Arrowheads in the final 3-6 frames point to micronuclei biogenesis events. Scale bars, 5µm. **(C)** The % of completed mitoses yielding micronuclei was calculated by dividing the number of mitoses yielding at least one micronucleus by the total number of completed mitoses captured during live imaging (Flox n=63; iKO n=52; pooled from 13 Flox and 22 iKO experiments). **(D)** All completed mitoses of Flox and iKO cells were binned into categories based on the number of timepoints observed in metaphase (Flox n=63; iKO n=52). Black and grey bars represent Flox and iKO data, respectively. **(E)** The % of premature mitotic exits was calculated by dividing the number of cells that entered prophase and returned to interphase without completing anaphase by the total number of cells that entered prophase (Flox n=71; iKO n=70; pooled from 13 Flox and 22 iKO experiments). 10 of the 70 iKO cells (14.3%) formed micronuclei. **(F)** iKO cells were fixed at 7d and stained with a γH2AX antibody (green) and DAPI (DNA; blue). Movies for panels A and B appear in Supporting Information (S1-S7 Videos).

However, iKO cell mitoses often yielded micronuclei due to errors in chromatin segregation (Fig 8A). This was repeatedly observed when chromatin fragments near the metaphase plate were not properly distributed to daughters during anaphase (Fig 8A). Another common phenotype of iKO cells that entered mitosis was premature mitotic exit (Fig 8B). Such cells underwent a prolonged prometaphase or metaphase, failed to enter anaphase, and ultimately returned to interphase with nuclei containing twice their normal chromatin content (Fig 8B). Micronuclei also formed in some of these cells that failed to complete mitosis (Fig 8B). We next measured the frequencies with which the mitotic defects occurred. First, approximately 23% of completed mitoses in iKO cells yielded micronuclei compared to only about 6% of completed mitoses in Flox cells (Fig 8C). Second, after categorizing the stages of mitosis and determining the number of timelapse frames spent specifically in metaphase (judged by the presence of at least 95% of H2B-GFP fluorescence aligned compactly at the cell equator), we found that >80% of Flox cells that completed mitosis spent only one frame in metaphase, whereas only 40% of iKO cells proceeded through metaphase with this speed (Fig 8D). The other 60% of iKO cells that completed mitosis spent two or more frames in metaphase (Fig 8D), suggesting that iKO cells experience an unusually prolonged metaphase period. Third, when evaluating the incidence of premature mitotic exits (defined as cells that entered prophase but did not complete anaphase), we discovered that nearly 30% of iKO cells that entered prophase underwent premature mitotic exits compared to only 10% of Flox cells (Fig 8E). Approximately 14% of premature mitotic exits that took place in the iKO cells also gave rise to micronuclei. Fourth, among the 330 iKO interphase nuclei that were observed during live cell imaging, at most 2 (i.e., ≤0.6%) appeared to create micronuclei via budding or extrusion, indicating that micronuclei form almost exclusively during mitosis. As a final point, to confirm that the prevalent chromatin missegregation events in iKO cells were, at least in part, attributable to the presence of broken DNA fragments, we fixed and stained mitotic cells for γH2AX. In agreement with this possibility, during prometaphase and metaphase, iKO cells contained γH2AX foci at the periphery of or completely detached from the main chromatin mass (Fig 8F). Furthermore, in early anaphase, iKO cells displayed prominent DNA damage clusters near their equators, and by late anaphase, γH2AX-positive lagging chromosomes became apparent (Fig 8F). Overall, these live and fixed cell studies show that, during Arp2/3-depleted conditions, damaged chromatin fragments persist and are incorporated into micronuclei as a result of defects in mitotic chromosome segregation.

### Actin filament penetration into the central spindle is diminished in Arp2/3-depleted cells

Upon discovering that knocking out the Arp2/3 complex leads to chromatin partitioning defects, we wanted to also assess how the loss of this actin nucleator might alter the actin and microtubule cytoskeletons during mitosis. Actin has been observed in various parts of meiotic and mitotic spindles in diverse organisms [30–39], so to examine the organization and intensity of actin filaments in relation to microtubule spindles in MTFs, we stained mitotic Flox and iKO cells with phalloidin to label F-actin, an anti-tubulin antibody to visualize microtubules, and DAPI to detect DNA. We focused on metaphase, when the chromosomes are either properly or improperly aligned at the central spindle, corresponding to when Flox mitoses rapidly proceed or iKO mitoses frequently stall. Imaging of metaphase Flox cells in multiple focal planes revealed that several distinct F-actin structures were present. In the lower portions of cells, linear actin filaments in the shape of a spindle and several bundles of microtubules were observed (Fig 9A). In the middle planes of cells, multiple thick finger-like F-actin structures penetrated the chromosomal region, and numerous microtubules comprising the main microtubule spindle were apparent (Fig 9A). In the upper parts of cells, fewer F-actin and microtubule structures intercalated the central spindle area (Fig 9A). Staining with anti-ArpC2 antibodies indicated that the Arp2/3 complex was not enriched along the thin linear actin filaments at the lower part of the cell, but was present near F-actin and microtubules in the middle and upper spindle structures (Fig 9A). Together, these observations expand the catalog of F-actin and Arp2/3-associated cytoskeletal structures that are found within dividing mammalian cells.

**Fig 9.**
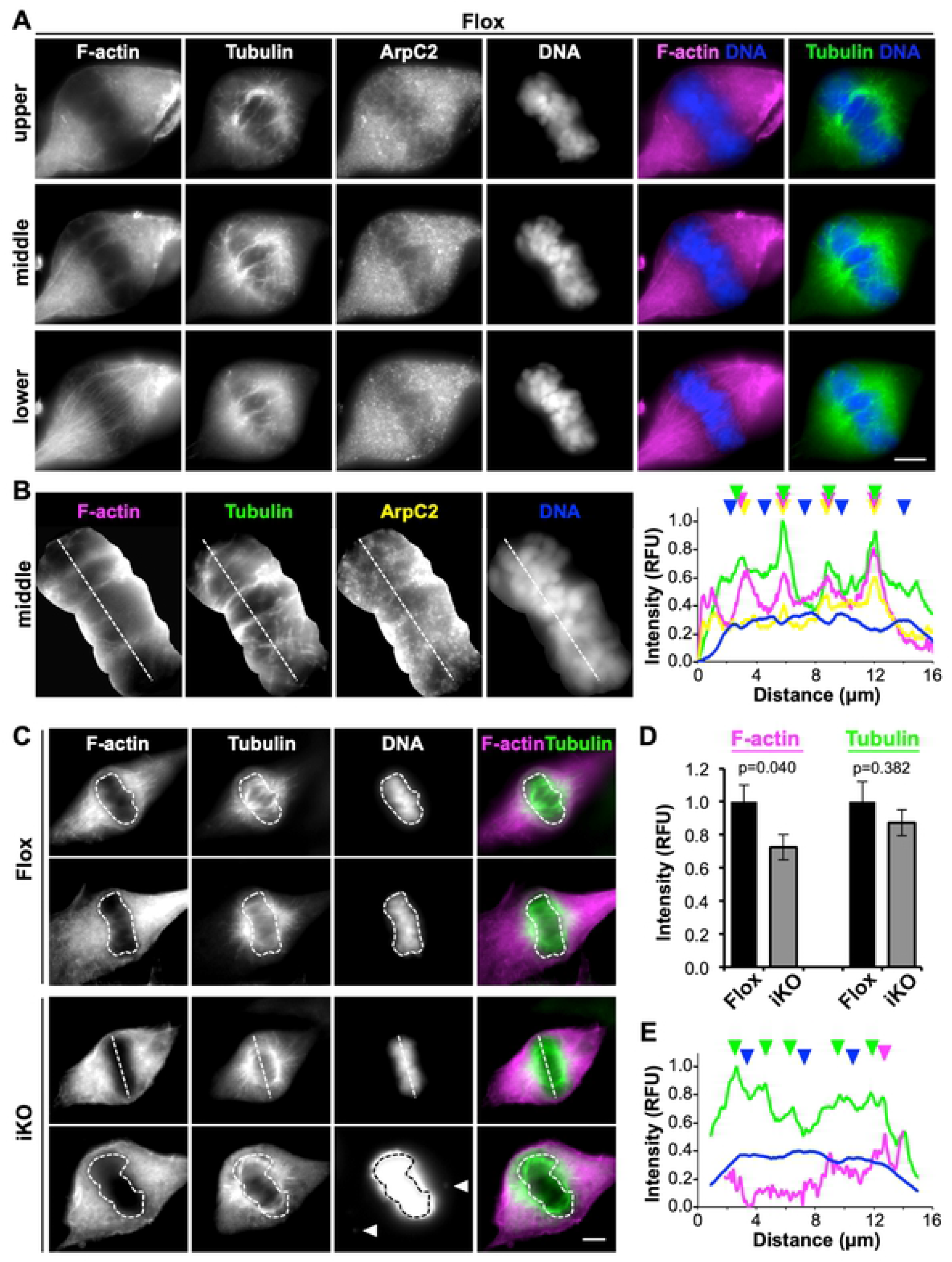
Arp2/3 complex depletion reduces actin filament density in the chromatin-containing central spindle during metaphase. **(A)** Mouse fibroblasts were treated with DMSO for 1-2d, fixed, and stained with phalloidin (F-actin; magenta), an anti-tubulin antibody (microtubules; green), anti-ArpC2 antibodies (yellow), and DAPI (DNA; blue). Images represent the lower, middle, and upper regions of metaphase cells. Scale bars, 5µm. **(B)** The DNA-containing region was isolated from the middle spindle in A, magnified, and subjected to line-scan fluorescence intensity analyses. Plot profiles depict the pixel intensity values for tubulin (green), F-actin (magenta), ArpC2 (yellow), and DNA (blue) along the dashed lines drawn across the representative metaphase region. Color-coded arrowheads highlight the overlapping peaks of tubulin, F-actin, and ArpC2 and the inverse intensity pattern for DNA. RFU = Relative Fluorescence Units. **(C)** Mouse fibroblasts were treated with DMSO (Flox) or 4-OHT (iKO) for 1-2d, fixed, and stained with phalloidin, an anti-tubulin antibody, and DAPI. The chromatin mass at the central metaphase spindle was outlined in ImageJ (dashed shapes) or a plot-profile line was drawn through it (dashed line). The bottom DAPI-stained iKO cell panel was overexposed in order to draw attention to the presence of 2 chromatin fragments erroneously missing from the metaphase plate (arrowheads). **(D)** Fluorescence intensities of F-actin and microtubules were measured in chromatin areas outlined as in panel C. Each bar represents the mean intensity ±SD from n=12 metaphase chromatin regions compiled from 3 experiments. **(E)** Plot profiles performed as in panel B depict the pixel intensity values for tubulin, F-actin, and DNA along the dashed lines drawn across the metaphase region from the representative iKO cell in panel C.

For more closely examining the spatial positioning of actin filaments, microtubules, and the Arp2/3 complex at the metaphase plate, we performed fluorescence intensity line scan analyses through the chromatin-containing region. Peaks of spindle microtubule intensity often coincided with peaks of F-actin intensity as well as sites of ArpC2 enrichment (Fig 9B). In contrast, DNA staining levels were highest where microtubules and F-actin were lowest (Fig 9B). These observations show a positive relationship between F-actin and microtubule localization in the central metaphase spindle.

To determine whether loss of the Arp2/3 complex causes any abnormalities in F-actin organization during metaphase, we compared F-actin staining in Flox and iKO cells. Actin filament levels in the metaphase chromatin-containing region appeared less prominent in iKO cells than in Flox cells (Fig 9C). Quantification of phalloidin fluorescence intensities in these regions of the central spindle revealed that F-actin levels were significantly lower in iKO cells compared to Flox cells (Fig 9D). This was not due to a general deficit in central spindle staining, because the fluorescence intensity of microtubules in the same region was not significantly different between iKO and Flox cells (Fig 9D). A reduction in the penetration of actin filaments, but not microtubules, into the metaphase chromatin-containing region of iKO cells was further reflected by fewer prominent F-actin peaks in fluorescence intensity plot profiles (Fig 9E). Collectively, these results indicate that a functional Arp2/3 complex is required for the polymerization of actin filaments in the vicinity of metaphase chromosomes.

### Arp2/3 depletion results in anaphase microtubule asymmetry

Since metaphase spindle-associated F-actin overlapped with microtubules in Flox cells and was less prominent in Arp2/3 knockout cells, we considered that the subsequent arrangement of microtubules during chromosome segregation might be altered. To explore this possibility, we evaluated microtubule organization and intensity during anaphase, when the chromosomes are either correctly or incorrectly partitioned. Flox cells exhibited uniform distributions of microtubules across the width of the separating spindle (Fig 10A), as evidenced by the relatively evenly-spaced peaks in fluorescence intensity plots perpendicular to the presumed position of the cytokinetic ring (Fig 10B). In contrast, iKO cells displayed unbalanced tubulin intensities in which the microtubules appeared to be more heavily concentrated on one side of the spindle (Fig 10A and 10B). These results suggest that in addition to the presence of misplaced damaged chromatin and a decrease in actin filaments at the metaphase plate, chromosome missegregations arising from alterations in anaphase microtubule organization may be a contributing factor in the formation of micronuclei in Arp2/3-deficient cells. Thus, loss of the Arp2/3 complex has multiple significant molecular consequences – beginning with unrepaired DNA damage and including mitotic actin and microtubule cytoskeleton abnormalities – that lead to a p53/p21 response, the biogenesis of micronuclei, cGAS/STING recruitment, and the onset of senescence.

**Fig 10.**
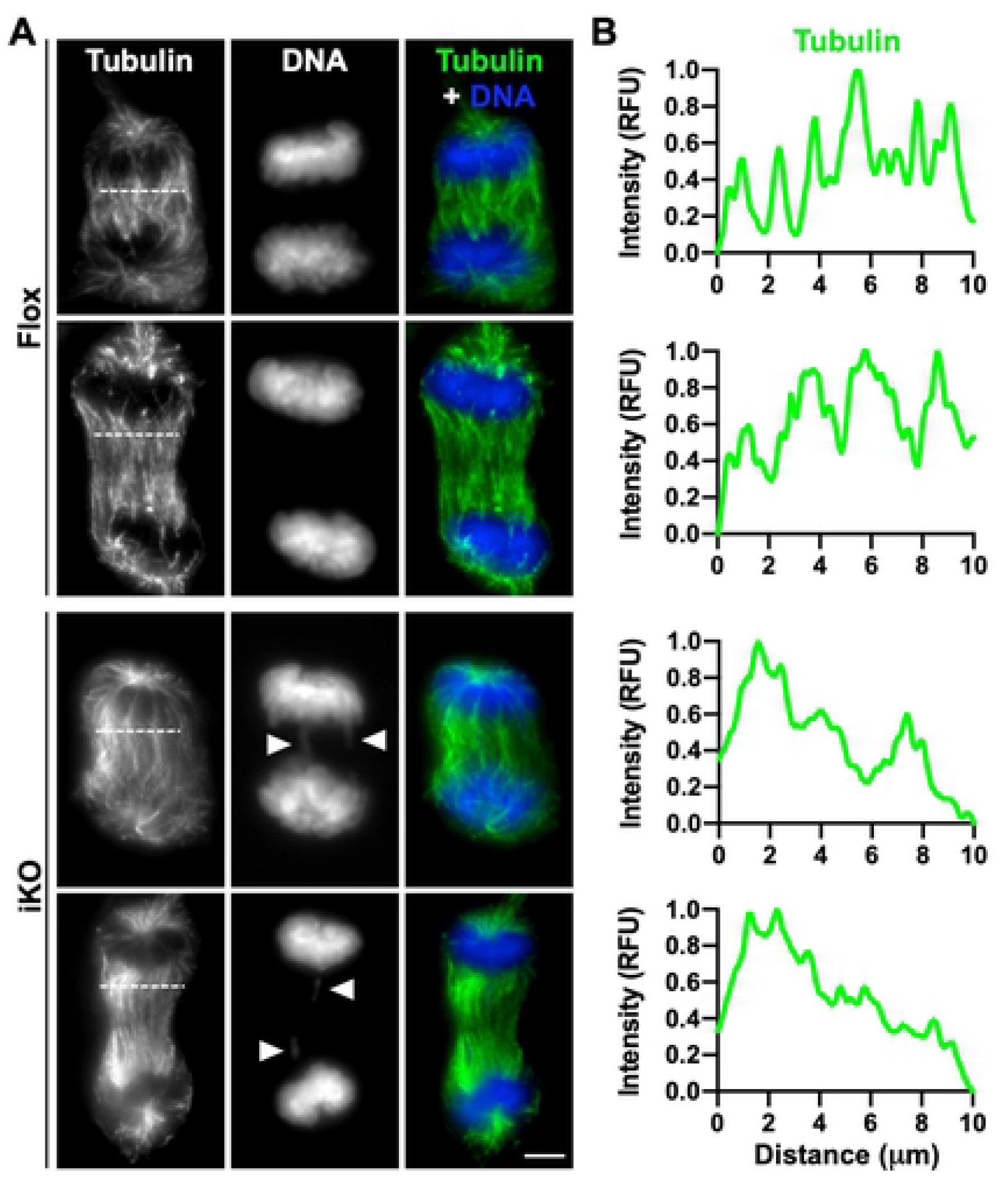
Anaphase microtubules are unbalanced in ArpC2 iKO cells. **(A)** Mouse fibroblasts were treated with DMSO (Flox) or 4-OHT (iKO) for 1d, fixed, and stained with a tubulin antibody (green) and DAPI (DNA; blue). Scale bar, 5µm. **(B)** Line-scan plot profiles depict the pixel intensity of tubulin along the horizontal dashed lines drawn across the representative mitotic spindles from panel A. Arrowheads highlight sites of aberrant chromatin localization. RFU = Relative Fluorescence Units.

## Discussion

The Arp2/3 complex is a key driver of many cellular processes that are mediated by actin assembly at the plasma membrane, namely adhesion, endocytosis, protrusion, and migration. While the complex is evolutionarily conserved among nearly all eukaryotes and is essential for viability in animals, the molecular basis underlying its indispensability is unclear. Roles for the Arp2/3 complex in enabling DNA repair during interphase, promoting chromosome partitioning during meiosis/mitosis, and controlling apoptosis following DNA damage have recently emerged and may help explain why Arp2/3 is essential. Such studies prompted us to determine the effects of permanent Arp2/3 complex ablation on viability, proliferation, and chromatin-associated processes in mammalian cells. Our results reveal important functions for the Arp2/3 complex in maintaining genomic integrity and supporting mitotic progression, while uncovering striking mechanistic connections between actin cytoskeleton defects and senescence.

Various endogenous stresses are known to induce cellular senescence, including replicative, telomeric, genotoxic, oncogenic, oxidative, and mitochondrial stress [81]. Our studies now add cytoskeletal dysfunction as a trigger of senescence. Arp2/3 complex knockout cells, in addition to a stable proliferation arrest and physical enlargement, display multiple phenotypic hallmarks of senescence. These include an increase in *Il-6* transcript reflective of SASP expression and a reduction of nuclear Lamin B1 levels. The iKO cells also harbor high cytoplasmic SA-βgal activity and increased acidic organelle content. Given that the Arp2/3 complex influences most cellular functions, from plasma membrane remodeling [2,15,82] to mitochondrial dynamics [83] to autophagy [84] to DNA repair [27, 28], it seems likely that multiple intracellular defects derived from Arp2/3 deficiency can impact senescence induction. Potential changes in mitochondrial turnover, combined with altered lysosomal content could signify increases in mitochondrial, oxidative, and proteotoxic stress in iKO cells. The extent to which these stressors contribute to the initiation or maintenance of senescence will be an important area of future investigation.

Despite uncertainties in the severity of cytoplasmic organelle dysfunction in iKO cells, obvious defects in nuclear and chromatin-associated processes emerged as major factors in promoting senescence in our studies. Kinetic analyses indicate that 1-3 days after exposure to 4-OHT, ArpC2 (and thus functional Arp2/3 complex) levels are 50-90% depleted (Fig 1A and 1B). This time period coincides with the steepest increases in detection of micronuclei via DAPI staining (Fig 4C) and visualization of dsDNA breaks in both nuclei and micronuclei via γH2AX staining (Fig 5C). A DNA damage response is activated, as evidenced by increased p53 expression and phosphorylation (Fig 6A and 6B). By 4-6 days, the p53-regulated cell cycle inhibitor p21 accumulates in the nucleus (Fig 6C-F) and cell multiplication stops (Fig 1D). By 7 days, SA-βgal-positivity begins to increase (Fig 3A and 3B). At 7-10 days, the entire population of iKO cells appears to be senescent, given the complete lack of proliferation (Fig 1D), as well as consistent single cell phenotypes in enlargement (Fig 1G), nuclear Lamin B1 reduction (Fig 2F), and cytoplasmic acidity (Fig 3D). Beyond 10 days, the iKO cell population remains viable but non-replicative while maintaining 20-25% positivity for micronuclei and SA-βgal activity.

While it is clear that all Arp2/3 knockout cells senesce, it is unclear why only a fraction of the culture is SA-βgal positive. This could be a technical matter of assay sensitivity, or intense SA-βgal-positivity may simply develop later in some cells. Additionally, the reasons why the entire culture senesces when only a quarter of the iKO cells harbor detectable micronuclei remain ambiguous. It is conceivable that aneuploidy as a result of receiving insufficient chromatin following mitosis or from excess DNA content following premature mitotic exits contributes to a p53-mediated arrest program in micronucleus-negative cells. Another possibility is that paracrine signaling from mature senescent cells to others in the culture supports the global proliferation block.

In response to damaging stimuli, the cyclin-dependent kinase inhibitors p16 and p21 are frequently upregulated and act as key players in pathways that promote the replicative arrest that defines senescence [55–57]. Earlier studies showed that inactivation of *p16Ink4a/Arf* is necessary for establishing Arp2/3-depleted mouse fibroblast cultures, suggesting that the loss of Arp2/3 can induce growth arrest or death in a p16-mediated manner [40, 41]. More recent experiments in human cells treated with a genotoxic agent indicate that depletion of the Arp2/3 complex or inactivation of two of its upstream regulators prevents the execution of apoptosis [29]. Such cells undergo a p53- and p21-associated cell cycle arrest but fail to properly complete an apoptosome-based caspase cleavage cascade [29]. These observations, together with our data showing that ArpC2 iKO cells activate p53 and accumulate p21 in a p16- independent manner, suggest that losing the Arp2/3 complex can trigger multiple cell cycle arrest pathways. While p53 is known to cause upregulation of *Cdkn1a*/p21 to promote arrest [85–87], deciphering the degree to which other anti-proliferative mechanisms cooperate with p21 in iKO cells requires more investigation. Interestingly, studies in aged mice show that cells expressing high levels of p21 are mostly distinct from those expressing high levels of p16 [65], further highlighting the possibility that loss of the Arp2/3 complex *in vivo* may instigate different pathways to senescence depending on the physiological state of the cell and its tissue of origin.

In addition to the functions of p53 and p21 in the nucleus, it seems likely that recognition of micronuclei in the cytoplasm also participates in the senescence response in Arp2/3-deficient cells. We observed cGAS and STING localization at or around micronuclei in ArpC2 iKO cells, and a chemical inhibitor of cGAS seemed to slightly increase multiplication of iKO cultures.

Recent work has also implicated cGAS/STING in regulating p21 expression, as depletion of cGAS or STING results in reduced p21 levels and premature mitotic entry [88]. These results support the notion that multiple nuclear and cytoplasmic signaling factors contribute to the induction and maintenance of the proliferation arrest in ArpC2 iKO cells.

The main mechanism underlying micronucleus biogenesis in Arp2/3-deficient cells is a lack of fidelity in chromatin partitioning during mitosis, as revealed by our live imaging studies.

Such segregation defects can be primarily explained by broken DNA fragments failing to properly attach to the microtubule spindle, since most micronuclei formed from cells completing inaccurate mitoses. However, cytoskeletal abnormalities may also contribute to chromosome segregation errors, as changes in the organization of metaphase F-actin and anaphase microtubules are also prevalent in the knockout cells. Our findings complement previous experiments showing altered F-actin levels in meiotic/mitotic structures and defects in spindle formation following chemical inhibition of Arp2/3 [37–39]. Interestingly, many mitoses in ArpC2 iKO cells feature a prolonged metaphase period. Delayed mitoses arising from failure to satisfy spindle assembly checkpoints can cause chromosome segregation mistakes and premature mitotic exits, both of which give rise to aneuploid cells [89], and both of which were observed at high frequencies in iKO cells. Thus, Arp2/3 complex deficiency leads to multiple deleterious consequences in M-phase of the cell cycle. More work is needed to determine how Arp2/3, actin, and microtubules collaborate in controlling spindle positioning, chromosome alignment, and DNA segregation.

Given that high mutation rates and aneuploidy are associated with organismal aging and tumorigenesis [90], impairment of Arp2/3 function *in vivo* could be a contributor to the development of age-related dysfunction and cancers. One central feature of the aging process is an accumulation of senescent cells, which can drive several aging phenotypes [91, 92]. Such discoveries have led to the development of new “senolytic” classes of drugs to treat and slow the progression of age-related pathologies [93]. Our findings suggest that impaired Arp2/3 complex function, and potentially other actin cytoskeletal misregulation, may contribute to premature senescence and aging. Taken together with previous observations that F-actin integrity affects aging and lifespan in *C.elegans* [94, 95], preventing or correcting cytoskeletal defects may promote cellular longevity and help reduce the senescent cell burden during organismal aging.

In contrast to the negative impacts of cellular senescence on aging, the anti-proliferative effects of senescence can serve as a positive form of tumor suppression [96]. Several therapeutics that induce senescence in cancer cells have been developed to reduce and prevent metastatic growth [97]. The human *CDKN2A* locus, which encodes p16^INK4A^/Arf, is frequently inactivated or epigenetically suppressed in various types of cancers [98]. Thus, the *ArpC2* floxed *Cdkn2a* null cells used in our studies present a unique opportunity to study Arp2/3 complex function in a genetic background that is particularly relevant to preventing the growth of cancer cells. Enhancing our understanding of the connection between the actin cytoskeleton and cellular senescence will therefore provide insight into therapeutic strategies used to regulate cell proliferation, arrest, and death in the context of both age-related diseases and cancers.

## Materials and Methods

### Ethics statement

Research with biological materials was approved by the UConn Institutional Biosafety Committee. This study did not include research with human subjects or live animals.

### Mammalian cell culture

MTFs containing the floxed *ArpC2* allele (from James Bear, University of North Carolina) [42] were cultured in DMEM (with 4.5g/L glucose, L-Glutamine, 110mg/L sodium pyruvate), 10% fetal bovine serum (FBS), GlutaMAX, and antibiotic-antimycotic (Gibco). Cells were treated with media containing 0.01% DMSO or 2µM 4-OHT (Sigma) to obtain Flox or iKO populations. For treatments exceeding 3 days, culture supernatants were replaced with fresh media containing DMSO or 4-OHT on day 4. Cultures were returned to normal media after day 6. All experiments were performed using cells that had been in active culture for 2-12 trypsinized passages.

### Cell proliferation assays

MTFs were cultured in 12-well plates and cell titers were routinely determined using a hemocytometer. Cells were initially seeded at multiple concentrations ranging from 2x10^3^ to 2x10^4^ cells per well. After 5 days, confluent Flox samples were subcultured daily into multiple wells at concentrations of 1-2x10^4^ cells per well, while iKO samples were subcultured if/when they reached 95% confluency. All cultures were expanded into 6-well plates and 6cm dishes when necessary. Population doubling times were calculated based on initial and final cell titers every 24-48h using the equation [time x log(2)] / [log(final) - log(initial)]. Due to their continuous proliferation, the plotted values of cell numbers for Flox samples at days 8-12 were extrapolations based on doubling rates at those time points. For cGAS inhibitor experiments, MTF supernatants were replaced with fresh media containing 5µM RU.521 every 12h from days 4-6 after 4-OHT exposure and counted at 0, 24, and 48h of DMSO or RU.521 treatment.

### Transfections and fluorescent probes

For mCherry-cGAS cloning, mouse cGAS was amplified via PCR from a pMSCVpuro-eGFP-cGAS template (Addgene, 108675) using primers containing KpnI and NotI restriction sites (S1 Table), and inserted into the pKC-mCherryC1 vector [99]. For transfections, Flox cells were grown in 12-well plates for 24h and then transfected with 350ng of mCherry-cGAS plasmid in DMEM. After 3h, DMEM was replaced with MTF media, and 18h later cells were trypsinized and transferred onto 12mm glass coverslips in 24-well plates. Media containing DMSO or 4-OHT was added after 3h, and cells were subjected to additional 29h growth before fixation as described below. For H2B-GFP transfections, Flox cells were grown in 24-well plates for 24h and then transfected with 130ng of H2B-GFP plasmid (Addgene, 11680). After 5h, DMEM was replaced with MTF media containing DMSO or 4-OHT. Cells were imaged live 15-40h later, as described below. All plasmids were maintained in NEB5-alpha *E.coli* and purified using Macherey-Nagel kits. For imaging acidic cytoplasmic organelles, cells were incubated for 30min in media containing 100nM LysoTracker Red (Invitrogen) prior to fixation.

### RT-PCR and RT-qPCR

RNA from Flox and iKO fibroblasts grown in 6-well plates was isolated using TRIzol reagent (Ambion). Following chloroform extraction, isopropanol precipitation, and a 75% ethanol wash, total RNA was resuspended in water. cDNA was reverse transcribed using the iScript cDNA synthesis kit (Bio-Rad) and then PCR-amplified using Taq polymerase (New England Biolabs) and primers listed in S1 Table. Primers were designed to amplify ∼340-480bp with each cDNA target. The resulting PCR products were visualized on ethidium-bromide stained agarose gels. RT-qPCR was performed using SYBR-green (Bio-Rad) on a CFX96 Real-Time System (Bio-Rad). 1µl of cDNA was used in each 10µl RT-qPCR reaction, and all samples were run in duplicate. Primer dilution curves were analyzed to ensure primer specificity. Ct values were normalized to GAPDH and/or actin. The iKO:Flox ratio (fold difference) at each timepoint was calculated by the comparative ΔΔ cycle threshold method.

### Cell extracts

For preparation of whole cell extracts, fibroblasts cultured in 6-well plates were collected in phosphate buffered saline (PBS) containing 1mM EDTA, centrifuged at 2,900rpm for 5.5min at 4°C, and lysed in 25mM HEPES (pH 7.4), 100mM NaCl, 1% Triton-X-100, 1mM EDTA, 1mM Na_3_VO_4_, 1mM NaF, 1mM phenylmethylsulfonyl fluoride, and 10µg/ml each of aprotonin, leupeptin, pepstatin, and chymostatin on ice. Lysates were mixed with Laemmli sample buffer, boiled, and centrifuged prior to SDS-PAGE analyses.

### Immunoblotting

Cell extract samples were separated on 12% SDS-PAGE gels before transfer to nitrocellulose membranes (GE Healthcare). Membranes were blocked in PBS containing 5% milk before probing with primary antibodies at concentrations listed in S2 Table. Following overnight incubation at 4°C, membranes were washed and treated with IRDye-680/800- (LI-COR) or horseradish peroxidase-conjugated (GE Healthcare) secondary antibodies. Infrared and chemiluminescent bands were visualized using a LI-COR Odyssey Fc Imaging System. Band intensities were measured using LI-COR Image Studio software. Densitometries of proteins-of-interest were normalized to GAPDH, tubulin, and/or actin loading controls.

### Immunostaining

For immunofluorescence, fibroblasts cultured on glass coverslips in 24-well plates were washed with PBS and fixed using PBS containing 2.5% paraformaldehyde for 30min. Following PBS washes, cells were permeabilized using PBS containing 0.1% Triton X-100 for 2min, washed, and stained with primary antibodies in PBS containing 1% bovine serum albumin, 1% FBS, and 0.02% azide for 45-60min as described in S2 Table. Cells were washed and treated with Alexa Fluor 488-, 555-, or 647-conjugated goat anti-rabbit, anti-mouse, or anti-rat secondary antibodies, 1µg/mL DAPI, and/or 0.2U/mL Alexa Fluor 488- or 647-labeled phalloidin (Invitrogen) for 35-45min as detailed in S2 Table. Following washes, coverslips were mounted on glass slides in Prolong Gold anti-fade (Invitrogen).

### Fluorescence microscopy

All fixed and live cells were imaged using a Nikon Eclipse Ti microscope equipped with Plan Apo 100X (1.45 NA), Plan Apo 60X (1.40 NA), or Plan Fluor 20X (0.5 NA) objectives, an Andor Clara-E camera, and a computer running NIS Elements Software. Most images were taken as single epifluorescence slices, whereas 60X images of mCherry-transfected cells (Fig 7) were taken as z-stacks with a 0.3µm step size, and fixed mitotic cells were taken with a 0.3µm (Fig 9) or 0.5µm (Fig 8, Fig 10) step size. Live cell imaging was performed in a 35°C chamber (Okolab). During live imaging, cells were cultured in fresh media containing 25mM HEPES (pH 7.4) and DMSO or 4-OHT. Images were captured using the 20x objective at 12min intervals. All image processing and analysis was conducted using ImageJ/FIJI software [100].

### Fluorescence quantification

Microscopic quantifications were performed using ImageJ. For fibroblast area calculations, cell perimeters were outlined based on phalloidin staining. For Lamin B1 abundance, the mean fluorescence intensity (MFI) of Lamin B1 staining in the nuclear area of the cell was measured and the background signal from outside the cell was subtracted from the MFI values. For normalization purposes, all MFI values for individual Flox and iKO cells were divided by the overall average MFI across 2 replicates for Flox cells. For acidic organelle analyses, the number of cells with diffuse LysoTracker staining was scored as positive or negative and divided by the total number of cells analyzed. To determine the LysoTracker area as a % of the total cell area, cell perimeters were outlined based on phalloidin staining and the Threshold tool was used to quantify the % of this area containing LysoTracker staining. For micronuclei and γH2AX quantifications, the number of cells with DAPI-stained micronuclei and the number of cells with prominent nuclear γH2AX clusters were quantified and divided by the total number of cells analyzed. Clusters were defined as isolated and discrete regions of γH2AX staining. For analyses of p21, the MFI of p21 staining in the nuclear area of the cell was measured and the average p21 MFI across 3 replicates for Flox and iKO cells was calculated. The average MFI of Flox cells was set to 1. For area-based assays of F-actin and microtubule intensities during metaphase, the DAPI-stained DNA mass was outlined in ImageJ and the MFI of phalloidin and anti-tubulin staining in this area was measured relative to DAPI. In linescan analyses of F-actin, tubulin, ArpC2, and DNA staining intensities in metaphase and/or anaphase cells, the ImageJ plot-profile tool was used. For metaphase, 14-16µm lines were drawn across the chromatin mass. For anaphase, 10-15µm lines were drawn across the mitotic spindle halfway between one set of chromosomes and the equator of the two forming daughter cells. In each plot, the minimum pixel intensity recorded along the line was subtracted from all values along the line to set the minimum to 0, and then all values were divided by the maximum to set the highest peak to 1. The numbers of cells analyzed in each type of assay are listed in the Figure Legends.

### β-galactosidase assays

SA-βgal activity was assessed using the Senescence β-Galactosidase Staining Kit (Cell Signaling Technologies, 9860). Fibroblasts were cultured in 6-well plates, washed once with PBS, fixed for 20min, and washed twice with PBS before incubation in SA-βgal staining solution at 37°C in the dark for 20-24h. Images were captured using an iPhone 7 on a bright-field microscope equipped with a 10x objective. The % of SA-βgal-positive cells was quantified by counting the number of intensely blue-colored cells divided by the total number of cells.

Samples were coded and scored in a blinded fashion.

### Reproducibility and Statistics

All conclusions were based on observations made from at least 4 separate experiments, while quantifications were based on data from 2-5 representative experiments. Statistical analyses were performed using GraphPad Prism software as noted in the Figure Legends. P-values for data sets including 2 conditions were determined using unpaired t-tests unless otherwise noted. Analyses of data sets involving +/- scoring used Fisher’s exact test. P-values <0.05 were considered statistically significant.

## Acknowledgements

We thank Jim Bear (UNC Chapel Hill) for providing cells harboring the floxed *ArpC2* allele, Ming Xu (UConn Health) for comments on senescence experiments, Tom Maresca (UMass Amherst) for input on mitosis experiments, Tom Perrotta for support with writing, and Campellone Lab members for their suggestions related to this manuscript.

## Supporting Information Captions

S1 Table. Primers.

S2 Table. Immunofluorescence and immunoblotting reagents.

S1 Video. Timelapse movie of a H2B-GFP-expressing Flox cell (see Fig 8A first row).

S2 Video. Timelapse movie of a H2B-GFP-expressing Flox cell (see Fig 8A second row).

S3 Video. Timelapse movie of a H2B-GFP-expressing iKO cell (see Fig 8A third row).

S4 Video. Timelapse movie of a H2B-GFP-expressing iKO cell (see Fig 8A fourth row).

S5 Video. Timelapse movie of a H2B-GFP-expressing iKO cell (see Fig 8B first row).

S6 Video. Timelapse movie of a H2B-GFP-expressing iKO cell (see Fig 8B second row).

S7 Video. Timelapse movie of a H2B-GFP-expressing iKO cell (see Fig 8B third row).

## References

1. Pollard TD. Actin and Actin-Binding Proteins. Cold Spring Harb Perspect Biol. 2016 Aug 1;8(8):a018226.

2. Campellone KG, Welch MD. A nucleator arms race: cellular control of actin assembly. Nat Rev Mol Cell Biol. 2010 Apr;11(4):237–51.

3. Welch MD, DePace AH, Verma S, Iwamatsu A, Mitchison TJ. The human Arp2/3 complex is composed of evolutionarily conserved subunits and is localized to cellular regions of dynamic actin filament assembly. J Cell Biol. 1997 Jul 28;138(2):375–84.

4. Abella JV, Galloni C, Pernier J, Barry DJ, Kjær S, Carlier MF, Way M. Isoform diversity in the Arp2/3 complex determines actin filament dynamics. Nat Cell Biol. 2016 Jan;18(1):76–86.

5. Schwob E, Martin RP. New yeast actin-like gene required late in the cell cycle. Nature. 1992 Jan 9;355(6356):179–82.

6. Winter DC, Choe EY, Li R. Genetic dissection of the budding yeast Arp2/3 complex: a comparison of the in vivo and structural roles of individual subunits. Proc Natl Acad Sci U S A. 1999 Jun 22;96(13):7288–93.

7. Zaki M, King J, Fütterer K, Insall RH. Replacement of the essential Dictyostelium Arp2 gene by its Entamoeba homologue using parasexual genetics. BMC Genet. 2007 Jun 6;8:28.

8. Hudson AM, Cooley L. A subset of dynamic actin rearrangements in Drosophila requires the Arp2/3 complex. J Cell Biol. 2002 Feb 18;156(4):677–87.

9. Stevenson V, Hudson A, Cooley L, Theurkauf WE. Arp2/3-dependent pseudocleavage [correction of psuedocleavage] furrow assembly in syncytial Drosophila embryos. Curr Biol. 2002 Apr 30;12(9):705–11.

10. Sawa M, Suetsugu S, Sugimoto A, Miki H, Yamamoto M, Takenawa T. Essential role of the C. elegans Arp2/3 complex in cell migration during ventral enclosure. J Cell Sci. 2003 Apr 15;116(Pt 8):1505–18.

11. Patel FB, Bernadskaya YY, Chen E, Jobanputra A, Pooladi Z, Freeman KL, Gally C, Mohler WA, Soto MC. The WAVE/SCAR complex promotes polarized cell movements and actin enrichment in epithelia during C. elegans embryogenesis. Dev Biol. 2008 Dec 15;324(2):297–309.

12. Yae K, Keng VW, Koike M, Yusa K, Kouno M, Uno Y, Kondoh G, Gotow T, Uchiyama Y, Horie K, Takeda J. Sleeping beauty transposon-based phenotypic analysis of mice: lack of Arpc3 results in defective trophoblast outgrowth. Mol Cell Biol. 2006 Aug;26(16):6185–96.

13. Vauti F, Prochnow BR, Freese E, Ramasamy SK, Ruiz P, Arnold HH. Arp3 is required during preimplantation development of the mouse embryo. FEBS Lett. 2007 Dec 11;581(29):5691–7.

14. Suraneni P, Rubinstein B, Unruh JR, Durnin M, Hanein D, Li R. The Arp2/3 complex is required for lamellipodia extension and directional fibroblast cell migration. J Cell Biol. 2012 Apr 16;197(2):239–51.

15. Rotty JD, Wu C, Bear JE. New insights into the regulation and cellular functions of the ARP2/3 complex. Nat Rev Mol Cell Biol. 2013 Jan;14(1):7–12.

16. Kim IH, Racz B, Wang H, Burianek L, Weinberg R, Yasuda R, Wetsel WC, Soderling SH. Disruption of Arp2/3 results in asymmetric structural plasticity of dendritic spines and progressive synaptic and behavioral abnormalities. J Neurosci. 2013 Apr 3;33(14):6081–92.

17. Zhou K, Muroyama A, Underwood J, Leylek R, Ray S, Soderling SH, Lechler T. Actin- related protein2/3 complex regulates tight junctions and terminal differentiation to promote epidermal barrier formation. Proc Natl Acad Sci U S A. 2013 Oct 1;110(40):E3820–9.

18. Zhou K, Sumigray KD, Lechler T. The Arp2/3 complex has essential roles in vesicle trafficking and transcytosis in the mammalian small intestine. Mol Biol Cell. 2015 Jun 1;26(11):1995–2004.

19. Wang PS, Chou FS, Ramachandran S, Xia S, Chen HY, Guo F, Suraneni P, Maher BJ, Li R. Crucial roles of the Arp2/3 complex during mammalian corticogenesis. Development. 2016 Aug 1;143(15):2741–52.

20. Papalazarou V, Swaminathan K, Jaber-Hijazi F, Spence H, Lahmann I, Nixon C, Salmeron- Sanchez M, Arnold HH, Rottner K, Machesky LM. The Arp2/3 complex is crucial for colonisation of the mouse skin by melanoblasts. Development. 2020 Nov 15;147(22):dev194555.

21. Mullins RD, Heuser JA, Pollard TD. The interaction of Arp2/3 complex with actin: nucleation, high affinity pointed end capping, and formation of branching networks of filaments. Proc Natl Acad Sci U S A. 1998 May 26;95(11):6181–6.

22. Machesky LM, Insall RH. Scar1 and the related Wiskott-Aldrich syndrome protein, WASP, regulate the actin cytoskeleton through the Arp2/3 complex. Curr Biol. 1998 Dec 17- 31;8(25):1347–56.

23. Steffen A, Faix J, Resch GP, Linkner J, Wehland J, Small JV, Rottner K, Stradal TE. Filopodia formation in the absence of functional WAVE- and Arp2/3-complexes. Mol Biol Cell. 2006 Jun;17(6):2581–91.

24. Nolen BJ, Tomasevic N, Russell A, Pierce DW, Jia Z, McCormick CD, Hartman J, Sakowicz R, Pollard TD. Characterization of two classes of small molecule inhibitors of Arp2/3 complex. Nature. 2009 Aug 20;460(7258):1031–4.

25. Hetrick B, Han MS, Helgeson LA, Nolen BJ. Small molecules CK-666 and CK-869 inhibit actin-related protein 2/3 complex by blocking an activating conformational change. Chem Biol. 2013 May 23;20(5):701–12.

26. Belin BJ, Lee T, Mullins RD. DNA damage induces nuclear actin filament assembly by Formin -2 and Spire-½ that promotes efficient DNA repair. [corrected]. Elife. 2015 Aug 19;4:e07735.

27. Caridi CP, D’Agostino C, Ryu T, Zapotoczny G, Delabaere L, Li X, Khodaverdian VY, Amaral N, Lin E, Rau AR, Chiolo I. Nuclear F-actin and myosins drive relocalization of heterochromatic breaks. Nature. 2018 Jul;559(7712):54–60.

28. Schrank BR, Aparicio T, Li Y, Chang W, Chait BT, Gundersen GG, Gottesman ME, Gautier J. Nuclear ARP2/3 drives DNA break clustering for homology-directed repair. Nature. 2018 Jul;559(7712):61–66.

29. King VL, Leclair NK, Coulter AM, Campellone KG. The actin nucleation factors JMY and WHAMM enable a rapid Arp2/3 complex-mediated intrinsic pathway of apoptosis. PLoS Genet. 2021 Apr 19;17(4):e1009512.

30. Lénárt P, Bacher CP, Daigle N, Hand AR, Eils R, Terasaki M, Ellenberg J. A contractile nuclear actin network drives chromosome congression in oocytes. Nature. 2005 Aug 11;436(7052):812–8.

31. Burdyniuk M, Callegari A, Mori M, Nédélec F, Lénárt P. F-Actin nucleated on chromosomes coordinates their capture by microtubules in oocyte meiosis. J Cell Biol. 2018 Aug 6;217(8):2661–2674.

32. Azoury J, Lee KW, Georget V, Rassinier P, Leader B, Verlhac MH. Spindle positioning in mouse oocytes relies on a dynamic meshwork of actin filaments. Curr Biol. 2008 Oct 14;18(19):1514–9.

33. Mogessie B, Schuh M. Actin protects mammalian eggs against chromosome segregation errors. Science. 2017 Aug 25;357(6353):eaal1647.

34. Kim HC, Jo YJ, Kim NH, Namgoong S. Small molecule inhibitor of formin homology 2 domains (SMIFH2) reveals the roles of the formin family of proteins in spindle assembly and asymmetric division in mouse oocytes. PLoS One. 2015 Apr 2;10(4):e0123438.

35. Kita AM, Swider ZT, Erofeev I, Halloran MC, Goryachev AB, Bement WM. Spindle-F-actin interactions in mitotic spindles in an intact vertebrate epithelium. Mol Biol Cell. 2019 Jul 1;30(14):1645–1654.

36. Farina F, Gaillard J, Guérin C, Couté Y, Sillibourne J, Blanchoin L, Théry M. The centrosome is an actin-organizing centre. Nat Cell Biol. 2016 Jan;18(1):65–75.

37. Plessner M, Knerr J, Grosse R. Centrosomal Actin Assembly Is Required for Proper Mitotic Spindle Formation and Chromosome Congression. iScience. 2019 May 31;15:274–281.

38. Farina F, Ramkumar N, Brown L, Samandar Eweis D, Anstatt J, Waring T, Bithell J, Scita G, Thery M, Blanchoin L, Zech T, Baum B. Local actin nucleation tunes centrosomal microtubule nucleation during passage through mitosis. EMBO J. 2019 Jun 3;38(11):e99843.

39. Inoue D, Obino D, Pineau J, Farina F, Gaillard J, Guerin C, Blanchoin L, Lennon-Duménil AM, Théry M. Actin filaments regulate microtubule growth at the centrosome. EMBO J. 2019 Jun 3;38(11):e99630.

40. Wu C, Asokan SB, Berginski ME, Haynes EM, Sharpless NE, Griffith JD, Gomez SM, Bear JE. Arp2/3 is critical for lamellipodia and response to extracellular matrix cues but is dispensable for chemotaxis. Cell. 2012 Mar 2;148(5):973–87.

41. Wu C, Haynes EM, Asokan SB, Simon JM, Sharpless NE, Baldwin AS, Davis IJ, Johnson GL, Bear JE. Loss of Arp2/3 induces an NF-κB-dependent, nonautonomous effect on chemotactic signaling. J Cell Biol. 2013 Dec 23;203(6):907–16.

42. Rotty JD, Wu C, Haynes EM, Suarez C, Winkelman JD, Johnson HE, Haugh JM, Kovar DR, Bear JE. Profilin-1 serves as a gatekeeper for actin assembly by Arp2/3-dependent and - independent pathways. Dev Cell. 2015 Jan 12;32(1):54–67.

43. Dimchev V, Lahmann I, Koestler SA, Kage F, Dimchev G, Steffen A, Stradal TEB, Vauti F, Arnold HH, Rottner K. Induced Arp2/3 Complex Depletion Increases FMNL2/3 Formin Expression and Filopodia Formation. Front Cell Dev Biol. 2021 Feb 1;9:634708.

44. Gournier H, Goley ED, Niederstrasser H, Trinh T, Welch MD. Reconstitution of human Arp2/3 complex reveals critical roles of individual subunits in complex structure and activity. Mol Cell. 2001 Nov;8(5):1041–52.

45. Hayflick L, Moorhead PS. The serial cultivation of human diploid cell strains. Exp Cell Res. 1961 Dec;25:585–621.

46. Hayflick L. The limited in vitro lifetime of human diploid cell strains. Exp Cell Res. 1965 Mar;37:614–36.

47. Coppé JP, Patil CK, Rodier F, Sun Y, Muñoz DP, Goldstein J, Nelson PS, Desprez PY, Campisi J. Senescence-associated secretory phenotypes reveal cell-nonautonomous functions of oncogenic RAS and the p53 tumor suppressor. PLoS Biol. 2008 Dec 2;6(12):2853–68.

48. Kuilman T, Michaloglou C, Vredeveld LC, Douma S, van Doorn R, Desmet CJ, Aarden LA, Mooi WJ, Peeper DS. Oncogene-induced senescence relayed by an interleukin-dependent inflammatory network. Cell. 2008 Jun 13;133(6):1019–31.

49. Coppé JP, Desprez PY, Krtolica A, Campisi J. The senescence-associated secretory phenotype: the dark side of tumor suppression. Annu Rev Pathol. 2010;5:99–118.

50. Basisty N, Kale A, Jeon OH, Kuehnemann C, Payne T, Rao C, Holtz A, Shah S, Sharma V, Ferrucci L, Campisi J, Schilling B. A proteomic atlas of senescence-associated secretomes for aging biomarker development. PLoS Biol. 2020 Jan 16;18(1):e3000599.

51. Freund A, Laberge RM, Demaria M, Campisi J. Lamin B1 loss is a senescence-associated biomarker. Mol Biol Cell. 2012;23(11):2066–2075.

52. Dimri GP, Lee X, Basile G, Acosta M, Scott G, Roskelley C, Medrano EE, Linskens M, Rubelj I, Pereira-Smith O, et al. A biomarker that identifies senescent human cells in culture and in aging skin in vivo. Proc Natl Acad Sci U S A. 1995 Sep 26;92(20):9363–7.

53. Lee BY, Han JA, Im JS, Morrone A, Johung K, Goodwin EC, Kleijer WJ, DiMaio D, Hwang ES. Senescence-associated beta-galactosidase is lysosomal beta-galactosidase. Aging Cell. 2006 Apr;5(2):187–95.

54. Kurz DJ, Decary S, Hong Y, Erusalimsky JD. Senescence-associated (beta)-galactosidase reflects an increase in lysosomal mass during replicative ageing of human endothelial cells. J Cell Sci. 2000 Oct;113 (Pt 20):3613–22.

55. Hernandez-Segura A, Nehme J, Demaria M. Hallmarks of Cellular Senescence. Trends Cell Biol. 2018 Jun;28(6):436–453.

56. DiMicco R, Krizhanovsky V, Baker D, d’Adda di Fagagna F. Cellular senescence in ageing: from mechanisms to therapeutic opportunities. Nat Rev Mol Cell Biol. 2021 Feb;22(2):75–95.

57. Kumari R, Jat P. Mechanisms of Cellular Senescence: Cell Cycle Arrest and Senescence Associated Secretory Phenotype. Front Cell Dev Biol. 2021 Mar 29;9:645593.

58. Ivanov A, Pawlikowski J, Manoharan I, van Tuyn J, Nelson DM, Rai TS, Shah PP, Hewitt G, Korolchuk VI, Passos JF, Wu H, Berger SL, Adams PD. Lysosome-mediated processing of chromatin in senescence. J Cell Biol. 2013 Jul 8;202(1):129–43.

59. Dou Z, Ghosh K, Vizioli MG, et al. Cytoplasmic chromatin triggers inflammation in senescence and cancer. Nature. 2017 Oct 19;550(7676):402–406.

60. Miller KN, Dasgupta N, Liu T, Adams PD, Vizioli MG. Cytoplasmic chromatin fragments- from mechanisms to therapeutic potential. Elife. 2021 Jan 29;10:e63728.

61. Rogakou EP, Pilch DR, Orr AH, Ivanova VS, Bonner WM. DNA double-stranded breaks induce histone H2AX phosphorylation on serine 139. J Biol Chem. 1998 Mar 6;273(10):5858–68.

62. Kastenhuber ER, Lowe SW. Putting p53 in Context. Cell. 2017 Sep;170(6):1062–78.

63. Hafner A, Bulyk ML, Jambhekar A, Lahav G. The multiple mechanisms that regulate p53 activity and cell fate. Nat Rev Mol Cell Biol. 2019 Apr;20(4):199–210.

64. Beauséjour CM, Krtolica A, Galimi F, Narita M, Lowe SW, Yaswen P, Campisi J. Reversal of human cellular senescence: roles of the p53 and p16 pathways. EMBO J. 2003 Aug 15;22(16):4212–22.

65. Wang, B., Wang, L., Gasek, N.S. et al. An inducible *p21*-Cre mouse model to monitor and manipulate *p21*-highly-expressing senescent cells in vivo. Nat Aging 1, 962–973 (2021).

66. Abbas T, Dutta A. p21 in cancer: intricate networks and multiple activities. Nat Rev Cancer. 2009 Jun;9(6):400–14.

67. El-Deiry WS. p21(WAF1) Mediates Cell-Cycle Inhibition, Relevant to Cancer Suppression and Therapy. Cancer Res. 2016 Sep 15;76(18):5189–91.

68. Chen Q, Sun L, Chen ZJ. Regulation and function of the cGAS-STING pathway of cytosolic DNA sensing. Nat Immunol. 2016 Sep 20;17(10):1142–9.

69. Hopfner KP, Hornung V. Molecular mechanisms and cellular functions of cGAS-STING signalling. Nat Rev Mol Cell Biol. 2020 Sep;21(9):501–521.

70. Glück S, Guey B, Gulen MF, Wolter K, Kang TW, Schmacke NA, Bridgeman A, Rehwinkel J, Zender L, Ablasser A. Innate immune sensing of cytosolic chromatin fragments through cGAS promotes senescence. Nat Cell Biol. 2017 Sep;19(9):1061–1070.

71. Mackenzie KJ, Carroll P, Martin CA, Murina O, Fluteau A, Simpson DJ, Olova N, Sutcliffe H, Rainger JK, Leitch A, Osborn RT, Wheeler AP, Nowotny M, Gilbert N, Chandra T, Reijns MAM, Jackson AP. cGAS surveillance of micronuclei links genome instability to innate immunity. Nature. 2017 Aug 24;548(7668):461-465.

72. Li T, Chen ZJ. The cGAS-cGAMP-STING pathway connects DNA damage to inflammation, senescence, and cancer. J Exp Med. 2018;215(5):1287–1299.

73. Volkman HE, Cambier S, Gray EE, Stetson DB. Tight nuclear tethering of cGAS is essential for preventing autoreactivity. Elife. 2019 Dec 6;8:e47491.

74. de Oliveira Mann CC, Hopfner KP. Nuclear cGAS: guard or prisoner? EMBO J. 2021 Aug 16;40(16):e108293.

75. Taguchi T, Mukai K, Takaya E, Shindo R. STING Operation at the ER/Golgi Interface. Front Immunol. 2021 May 3;12:646304.

76. Vincent J, Adura C, Gao P, Luz A, Lama L, Asano Y, Okamoto R, Imaeda T, Aida J, Rothamel K, Gogakos T, Steinberg J, Reasoner S, Aso K, Tuschl T, Patel DJ, Glickman JF, Ascano M. Small molecule inhibition of cGAS reduces interferon expression in primary macrophages from autoimmune mice. Nat Commun. 2017 Sep 29;8(1):750.

77. Fenech M, Kirsch-Volders M, Natarajan AT, Surralles J, Crott JW, Parry J, Norppa H, Eastmond DA, Tucker JD, Thomas P. Molecular mechanisms of micronucleus, nucleoplasmic bridge and nuclear bud formation in mammalian and human cells. Mutagenesis. 2011 Jan;26(1):125–32.

78. Utani K, Okamoto A, Shimizu N. Generation of micronuclei during interphase by coupling between cytoplasmic membrane blebbing and nuclear budding. PLoS One. 2011 Nov;6(11):e27233.

79. Crasta K, Ganem NJ, Dagher R, et al. DNA breaks and chromosome pulverization from errors in mitosis. Nature. 2012 Feb 2;482(7383):53–58.

80. Liu S, Pellman D. The coordination of nuclear envelope assembly and chromosome segregation in metazoans. Nucleus. 2020 Dec;11(1):35–52.

81. Gorgoulis V, Adams PD, Alimonti A, Bennett DC, Bischof O, Bishop C, Campisi J, Collado M, Evangelou K, Ferbeyre G, Gil J, Hara E, Krizhanovsky V, Jurk D, Maier AB, Narita M, Niedernhofer L, Passos JF, Robbins PD, Schmitt CA, Sedivy J, Vougas K, von Zglinicki T, Zhou D, Serrano M, Demaria M. Cellular Senescence: Defining a Path Forward. Cell. 2019 Oct;179(4):813–827.

82. Swaney KF, Li R. Function and regulation of the Arp2/3 complex during cell migration in diverse environments. Curr Opin Cell Biol. 2016;42:63–72.

83. Moore AS, Holzbaur ELF. Mitochondrial-cytoskeletal interactions: dynamic associations that facilitate network function and remodeling. Curr Opin Physiol. 2018 Jun;3:94–100.

84. Chakrabarti R, Lee M, Higgs HN. Multiple roles for actin in secretory and endocytic pathways. Curr Biol. 2021 May 24;31(10):R603–R618.

85. Basile JR, Eichten A, Zacny V, Münger K. NF-kappaB-mediated induction of p21(Cip1/Waf1) by tumor necrosis factor alpha induces growth arrest and cytoprotection in normal human keratinocytes. Mol Cancer Res. 2003 Feb;1(4):262–70.

86. Jackson JG, Pereira-Smith OM. p53 is preferentially recruited to the promoters of growth arrest genes p21 and GADD45 during replicative senescence of normal human fibroblasts. Cancer Res. 2006 Sep 1;66(17):8356–60.

87. Rovillain E, Mansfield L, Caetano C, Alvarez-Fernandez M, Caballero OL, Medema RH, Hummerich H, Jat PS. Activation of nuclear factor-kappa B signalling promotes cellular senescence. Oncogene. 2011 May 19;30(20):2356–66.

88. Basit A, Cho MG, Kim EY, Kwon D, Kang SJ, Lee JH. The cGAS/STING/TBK1/IRF3 innate immunity pathway maintains chromosomal stability through regulation of p21 levels. Exp Mol Med. 2020 Apr;52(4):643–657.

89. Rieder CL, Maiato H. Stuck in division or passing through: what happens when cells cannot satisfy the spindle assembly checkpoint. Dev Cell. 2004 Nov;7(5):637–51.

90. Collado M, Blasco MA, Serrano M. Cellular senescence in cancer and aging. Cell. 2007 Jul 27;130(2):223–33.

91. Baker DJ, Wijshake T, Tchkonia T, LeBrasseur NK, Childs BG, van de Sluis B, Kirkland JL, van Deursen JM. Clearance of p16Ink4a-positive senescent cells delays ageing-associated disorders. Nature. 2011 Nov 2;479(7372):232–6.

92. Baker DJ, Childs BG, Durik M, Wijers ME, Sieben CJ, Zhong J, Saltness RA, Jeganathan KB, Verzosa GC, Pezeshki A, Khazaie K, Miller JD, van Deursen JM. Naturally occurring p16(Ink4a)-positive cells shorten healthy lifespan. Nature. 2016 Feb 11;530(7589):184–9.

93. Hickson LJ, Langhi Prata LGP, Bobart SA, Evans TK, Giorgadze N, Hashmi SK, Herrmann SM, Jensen MD, Jia Q, Jordan KL, Kellogg TA, Khosla S, Koerber DM, Lagnado AB, Lawson DK, LeBrasseur NK, Lerman LO, McDonald KM, McKenzie TJ, Passos JF, Pignolo RJ, Pirtskhalava T, Saadiq IM, Schaefer KK, Textor SC, Victorelli SG, Volkman TL, Xue A, Wentworth MA, Wissler Gerdes EO, Zhu Y, Tchkonia T, Kirkland JL. Senolytics decrease senescent cells in humans: Preliminary report from a clinical trial of Dasatinib plus Quercetin in individuals with diabetic kidney disease. EBioMedicine. 2019 Sep;47:446–456.

94. Baird NA, Douglas PM, Simic MS, Grant AR, Moresco JJ, Wolff SC, Yates JR 3rd, Manning G, Dillin A. HSF-1-mediated cytoskeletal integrity determines thermotolerance and life span. Science. 2014 Oct 17;346(6207):360–3.

95. Higuchi-Sanabria R, Paul JW 3rd, Durieux J, Benitez C, Frankino PA, Tronnes SU, Garcia G, Daniele JR, Monshietehadi S, Dillin A. Spatial regulation of the actin cytoskeleton by HSF-1 during aging. Mol Biol Cell. 2018 Oct 15;29(21):2522–2527.

96. Hinds P, Pietruska J. Senescence and tumor suppression. F1000Res. 2017 Dec 11;6:2121.

97. Saleh T, Bloukh S, Carpenter VJ, Alwohoush E, Bakeer J, Darwish S, Azab B, Gewirtz DA. Therapy-Induced Senescence: An “Old” Friend Becomes the Enemy. Cancers (Basel). 2020 Mar 29;12(4):822.

98. Ko A, Han SY, Song J. Dynamics of ARF regulation that control senescence and cancer. BMB Rep. 2016 Nov;49(11):598–606.

99. Campellone KG, Webb NJ, Znameroski EA, Welch MD. WHAMM is an Arp2/3 complex activator that binds microtubules and functions in ER to Golgi transport. Cell. 2008 Jul;134(1):148–161.

100. Schindelin J, Arganda-Carreras I, Frise E, Kaynig V, Longair M, Pietzsch T, Preibisch S, Rueden C, Saalfeld S, Schmid B, Tinevez JY, White DJ, Hartenstein V, Eliceiri K, Tomancak P, Cardona A. Fiji: an open-source platform for biological-image analysis. Nat Methods. 2012 Jun 28;9(7):676–82.

